# eIF4E1b is a non-canonical eIF4E required for maternal mRNA dormancy

**DOI:** 10.1101/2023.06.10.544440

**Authors:** Laura Lorenzo-Orts, Marcus Strobl, Benjamin Steinmetz, Friederike Leesch, Carina Pribitzer, Michael Schutzbier, Gerhard Dürnberger, Andrea Pauli

## Abstract

Maternal mRNAs are essential for protein synthesis during oogenesis and early embryogenesis. To adapt translation to specific needs during development, maternal mRNAs are translationally repressed by shortening the polyA tails. While mRNA deadenylation is associated with decapping and degradation in somatic cells, maternal mRNAs with short polyA tails are stable. Here we report an essential role for the germline-specific paralog of the mRNA cap-binding factor eIF4E, known as eIF4E1b, in the storage and repression of maternal mRNAs with short polyA tails. eIF4E1b binds to the mRNA cap and is targeted to ribonucleoprotein complexes through its direct interaction with eIF4ENIF1/4E-T. In early embryos, eIF4E1b binds to a specific set of translationally repressed mRNAs with short or no polyA tails, such as histone mRNAs, which are translated later on during embryogenesis. Consistent with an important role in maternal mRNA dormancy, mutation of *eIF4E1b* in zebrafish impairs female germline development. Understanding the mechanism and function of eIF4E1B provides new insights into fundamental post-transcriptional regulatory principles governing early vertebrate development.

## Introduction

Eggs contain a large number of ribosomes and mRNAs that enable protein synthesis in the embryo. However, in order to maintain a state of quiescence, several mechanisms repress translation in the egg. Maternal mRNAs are translationally repressed by shortening of the polyadenine (polyA) tails (Subtelny *et al*, 2014; Lee *et al*, 2023). mRNAs with short polyA tails are not efficiently recognized by the polyA-binding protein (PABP) in the egg and thus cannot participate in polyA-dependent translation (Xiang & Bartel, 2021). In addition, recent work has shown that maternal ribosomes associate with several factors that block key functional sites of the ribosome and contribute to their dormancy (Leesch *et al*, 2023).

Since transcription stops at the onset of meiosis during oogenesis, eggs face the additional challenge of storing proteins and RNAs for later use in the early embryo. In zebrafish, for example, zygotic genome activation (ZGA) does not begin until 3 hours post fertilization (hpf), whereas in mouse it begins at approximately 24 hpf (Jukam *et al*, 2017; Vastenhouw *et al*, 2019). In contrast to somatic cells, where mRNA deadenylation leads to decapping and degradation (Passmore & Coller, 2022), mRNAs with short polyA tails are stable in the oocyte and early embryo (Voeltz & Steitz, 1998; Bhat *et al*, 2023; Lee *et al*, 2023). While several RNA-binding proteins, including Zar1, Zar1l/Zar2, MSY2/Ybx2, and Igf2bp3, which bind to untranslated regions (UTRs) or coding sequences of mRNAs, have been implicated in stabilizing maternal mRNAs, it is unclear how these proteins protect maternal transcripts with short polyA tails from decapping at the 5’ end (Gillian-Daniel *et al*, 1998; Rong *et al*, 2019; Medvedev *et al*, 2008, 2011; Ren *et al*, 2020).

In somatic cells, the mRNA cap-binding factor eIF4E plays a key role in regulating translation and mRNA stability. eIF4E interacts with the scaffolding protein eIF4G in the cytoplasm, which contributes to the formation of the heterotrimeric complex eIF4F, consisting of eIF4E, eIF4G and the RNA helicase eIF4A (Gingras *et al*, 1999). eIF4F is essential for canonical (cap- and polyA-dependent) translation and can be inhibited by eIF4EBPs, which compete with eIF4G for eIF4E binding (Marcotrigiano *et al*, 1999). eIF4E also interacts with the P-body component eIF4ENIF1/4E-T to inhibit mRNA decapping (Räsch *et al*, 2020). Decapping requires eIF4E to dissociate from the mRNA cap, which is otherwise inaccessible to the decapping enzyme Dcp2 (Vilela *et al*, 2000; Schwartz & Parker, 2000). eIF4E interactions depend on the presence of eIF4E-binding motifs (consisting of YXXXXLΦ; where X is any amino acid and Φ is a hydrophobic residue) in eIF4E-binding proteins, including eIF4EG, eIF4EBP, and eIF4ENIF1 (Mader *et al*, 1995; Dostie *et al*, 2000). Vertebrates have evolved specific eIF4E classes (i.e., eIF4E2/4EHP and eIF4E3) that perform unique functions, such as translational inhibition upon ribosome collision and translational initiation upon stress, respectively (Juszkiewicz *et al*, 2020; Weiss *et al*, 2021).

Here, we report that the germline eIF4E paralog eIF4E1B is involved in maternal mRNA dormancy by binding to the mRNA cap of mRNAs with short polyA tails, contributing to their stability and repression.

## Results

Given the function of eIF4Es in regulating translation and mRNA stability, we hypothesized that eIF4Es may also contribute to maternal mRNA dormancy. To explore this possibility, we analyzed the expression of zebrafish *eIF4Es* during oogenesis and embryogenesis. Vertebrates have evolved three eIF4E classes with different affinities for the mRNA cap (Joshi *et al*, 2005). Class I eIF4Es contain two conserved tryptophans that interact with the mRNA cap. In class II (eIF4E2/4EHP) and class III (eIF4E3), one of these tryptophans is substituted by a different amino acid (Phe, Leu or Tyr in the case of eIF4E2; Cys in the case of eIF4E3), which reduces the affinity for the mRNA cap (Zuberek *et al*, 2007; Osborne *et al*, 2013). Moreover, class II and class III eIF4Es contain other amino acid substitutions that affect their interaction with eIF4Gs (in the case of eIF4E2) and eIF4EBPs (in the case of eIF4E3) (Joshi *et al*, 2004). Zebrafish have seven eIF4Es, four of which belong to class I (eIF4Ea, eIF4Eb, eIF4E1c and eIF4E1b), two to class II (eIF4E2 and eIF4E2rs1) and one to class III (eIF4E3) (**Fig 1A and B, Fig EV1**). Zebrafish class I eIF4Es share between 65 and 84% sequence identity with human eIF4E (**Fig 1B**). eIF4Ea and eIF4Eb are the most similar to mammalian eIF4E and are likely the result of a gene duplication event in fish (**Fig 1A and B, Fig EV1**) (Taylor *et al*, 2001). While eIF4E1c is specific to fish (**Fig EV1**) (Rao *et al*, 2023), eIF4E1b proteins are conserved in most vertebrates (**Figs EV1 and EV2A**). Expression data (published polyA-selected RNA-seq and tandem mass tag mass spectrometry, TMT-MS) show that eIF4E1b and eIF4E1c are highly expressed during zebrafish oogenesis and early embryogenesis (**Fig 1C and D)** (Pauli *et al*, 2012; Cabrera-Quio *et al*, 2021). Consistent with previous reports from zebrafish (Robalino *et al*, 2004; Rao *et al*, 2023) and mouse (Evsikov *et al*, 2006; Yang *et al*, 2022; Guo *et al*, 2023), our analyses indicate that zebrafish eIF4E1b is restricted to the germline and to early embryonic development (**Fig 1C, Fig EV2B**).

**Figure 1.**
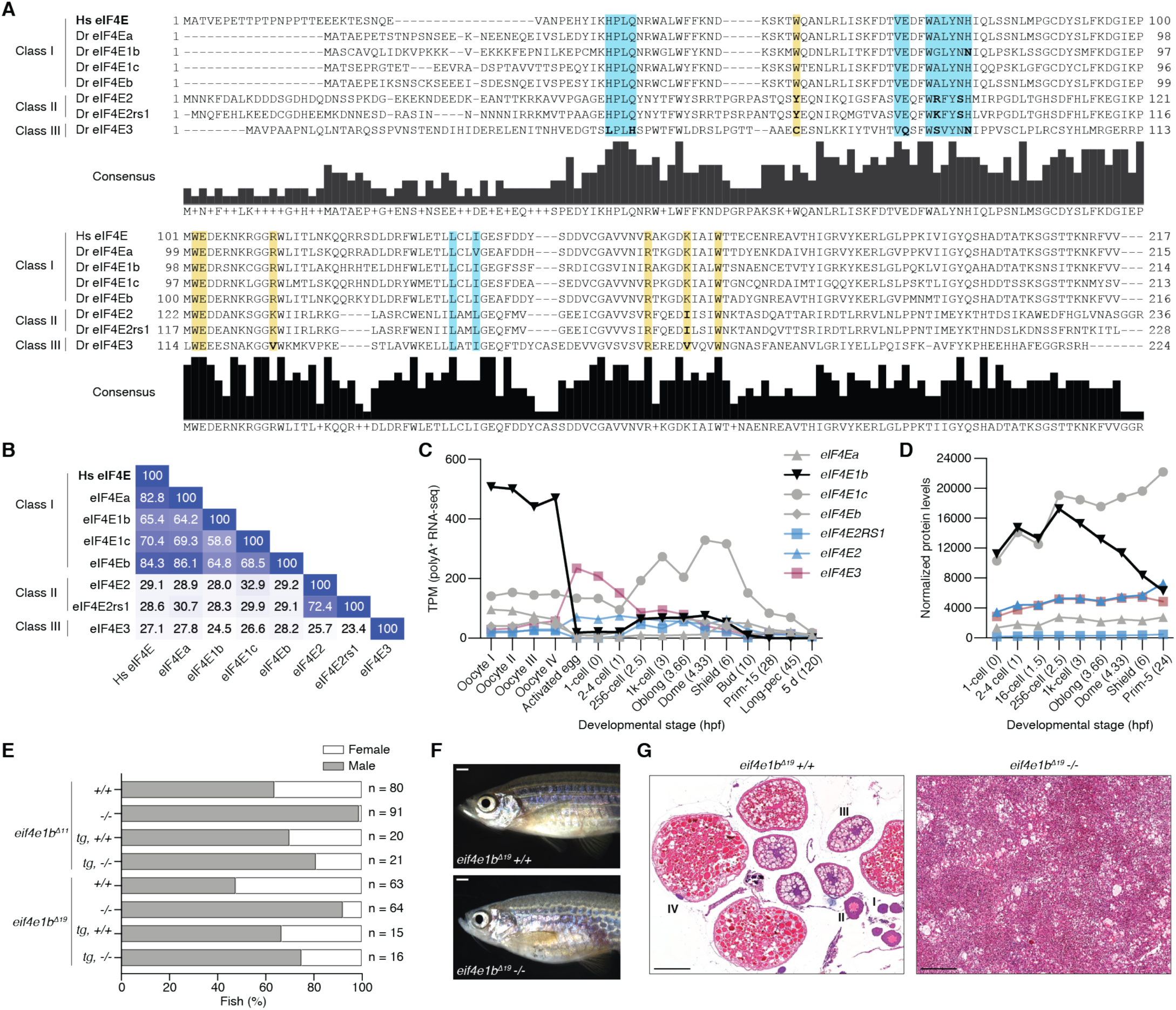
eIF4E1b is a class I eIF4E protein with an essential role in zebrafish oogenesis. **A** Alignment of human (*Homo sapiens*, Hs) and zebrafish (*Danio rerio*, Dr) eIF4E proteins. Highlighted regions indicate the amino acids involved in the interaction with the mRNA cap (yellow) and eIF4E-binding motifs (blue); residues in the highlighted regions with a different polarity than human eIF4E are indicated in bold. **B** Percentage identity matrix of the proteins shown in *A*. **C** Abundance of *eIF4E* transcripts during zebrafish oogenesis and embryogenesis based on published polyA-selected RNA-seq data (Pauli *et al*, 2012; Cabrera-Quio *et al*, 2021). TPM: transcripts per million. X-axis indicates developmental stages (hours post fertilization, hpf, in brackets). **D** eIF4E protein levels (represented with the same colors and symbols as in *C*) during early zebrafish embryogenesis obtained by tandem mass tag mass-spectrometry (TMT-MS), normalized to spike-in proteins. X-axis indicates developmental stages (hpf in brackets). **E** Percentage of males and females from homozygous (-/-) and wild-type (+/+) siblings obtained from heterozygous *eif4e1b* incrosses (n = number of fish), as determined by secondary sexual characteristics. Expression of 3xflag-sfGFP-eIF4E1b (*tg*) partially rescues the male bias observed in the two homozygous fish mutants. **F** Representative pictures of wild-type (top) and homozygous *eif4e1b* (bottom) female siblings (scale bar = 1 mm). **G** Hematoxylin and eosin staining of ovaries isolated from wild-type and homozygous *eif4e1b* fish reveals the absence of oocytes in mutant ovaries. Stages of oocyte development are indicated with roman numbers. Scale bars = 200 μm.

To understand the physiological relevance of eIF4E1b, we generated two CRISPR/Cas9-based *eif4e1b* knockout mutants in zebrafish containing different deletions in the third exon of the locus, resulting in frameshifts leading to premature stop codons (**Fig EV3A-C**). Most homozygous *eif4e1b* mutants developed into fertile males, and only a small proportion developed into infertile fish that morphologically resembled females but had gonads with tumor-like growth and no oocytes (**Fig 1E-G, Fig EV3D-G**). Importantly, ubiquitous expression of GFP-tagged eIF4E1b under the control of the *actb2* (*actin, beta 2*) promoter in homozygous *eif4e1b* fish rescued the defect in female development and resulted in fertile males and females (**Fig 1E, Fig EV3E and F**). Since oocytes are necessary to maintain a female sex in zebrafish (Dranow *et al*, 2013), the male bias observed in *eif4e1b* adults could be due to sex reversal. To investigate this, we used the *ziwi:eGFP* reporter to identify juveniles (1-2 cm long fish still lacking secondary sexual characteristics) that had started to develop as females based on a high GFP expression in the gonads (Leu & Draper, 2010; Dranow *et al*, 2016). While both homozygous *eif4e1b* mutant and wild-type siblings showed putative females with high GFP expression in their gonads as juveniles, only wild types developed into adult females (**Fig EV3H and I**), suggesting that loss of *eIF4E1b* causes sex reversal in zebrafish. In line with this, high GFP-expressing gonads from homozygous *eif4e1b* juveniles were either ovaries (with similar morphology as those isolated from wild types) or gonads containing differentiating sperm, the latter most likely representing ovaries in the process of transforming into testes (**Fig EV3J**). We therefore conclude that eIF4E1b is required for female germline development in zebrafish.

Although eIF4E1Bs belong to class I eIF4Es, they have been reported not to bind (Robalino *et al*, 2004) or to bind weakly (Minshall *et al*, 2007; Kubacka *et al*, 2015) to the mRNA cap, despite containing the two conserved tryptophan residues that are responsible for binding to the mRNA cap in canonical eIF4Es (**Figs 1A and 2A**). To test the ability of eIF4E1Bs to bind to the mRNA cap, we performed in vitro immunoprecipitation assays with m^7^G-coated beads and bacterial lysates containing soluble His and MBP tagged eIF4Es. As recombinant eIF4E1b from zebrafish was unstable in solution (**Fig EV4A**), we used mouse and human eIF4E1Bs for our in vitro studies (**Fig EV2A**). We observed that eIF4E1B binds to m^7^G with an affinity similar to that of eIF4Ea (**Fig 2B, Fig EV4B**), which is 83% identical to human eIF4E and contains all the residues involved in mRNA cap binding (**Fig 1A and B**). In support of the specificity of this interaction, the affinity of eIF4E1B for m^7^G was reduced when the tryptophans involved in mRNA cap binding in canonical eIF4E were mutated, suggesting that eIF4E1B binds to the mRNA cap in the same manner as eIF4E (**Fig 2A and B, Fig EV4B**). To investigate whether eIF4E1B can also bind to the eIF4E-binding motifs of eIF4G, eIF4EBP1 and eIF4ENIF1 like other class I eIF4Es, we performed in vitro pulldown experiments with bacterial lysates (**Fig 2C and D, Fig EV4C-E**). eIF4Ea and eIF4E1c bound to all tested eIF4E-binding motifs with similar affinities as murine eIF4E, although eIF4Ea showed a slightly higher affinity for eIF4ENIF1 (**Fig 2D**). In agreement with previous studies using mouse or human eIF4Es, we observed that zebrafish eIF4E3, but not eIF4E2, bind to eIF4G (**Fig 2D**) (Joshi *et al*, 2004; Osborne *et al*, 2013; Weiss *et al*, 2021). Unlike other class I eIF4Es, neither mouse nor human eIF4E1B interacted with eIF4G, although they bound to eIF4EBP1 and eIF4ENIF1 (**Fig 2D, Fig EV4F**). Taken together, our in vitro data show that, contrary to previous reports (Robalino *et al*, 2004; Minshall *et al*, 2007; Kubacka *et al*, 2015), eIF4E1Bs efficiently bind to the mRNA cap and to the eIF4E-binding motifs of eIF4EBP1 and eIF4ENIF1, yet do not interact with eIF4G and thus may not be promoting translation initiation like other class I eIF4Es.

**Figure 2.**
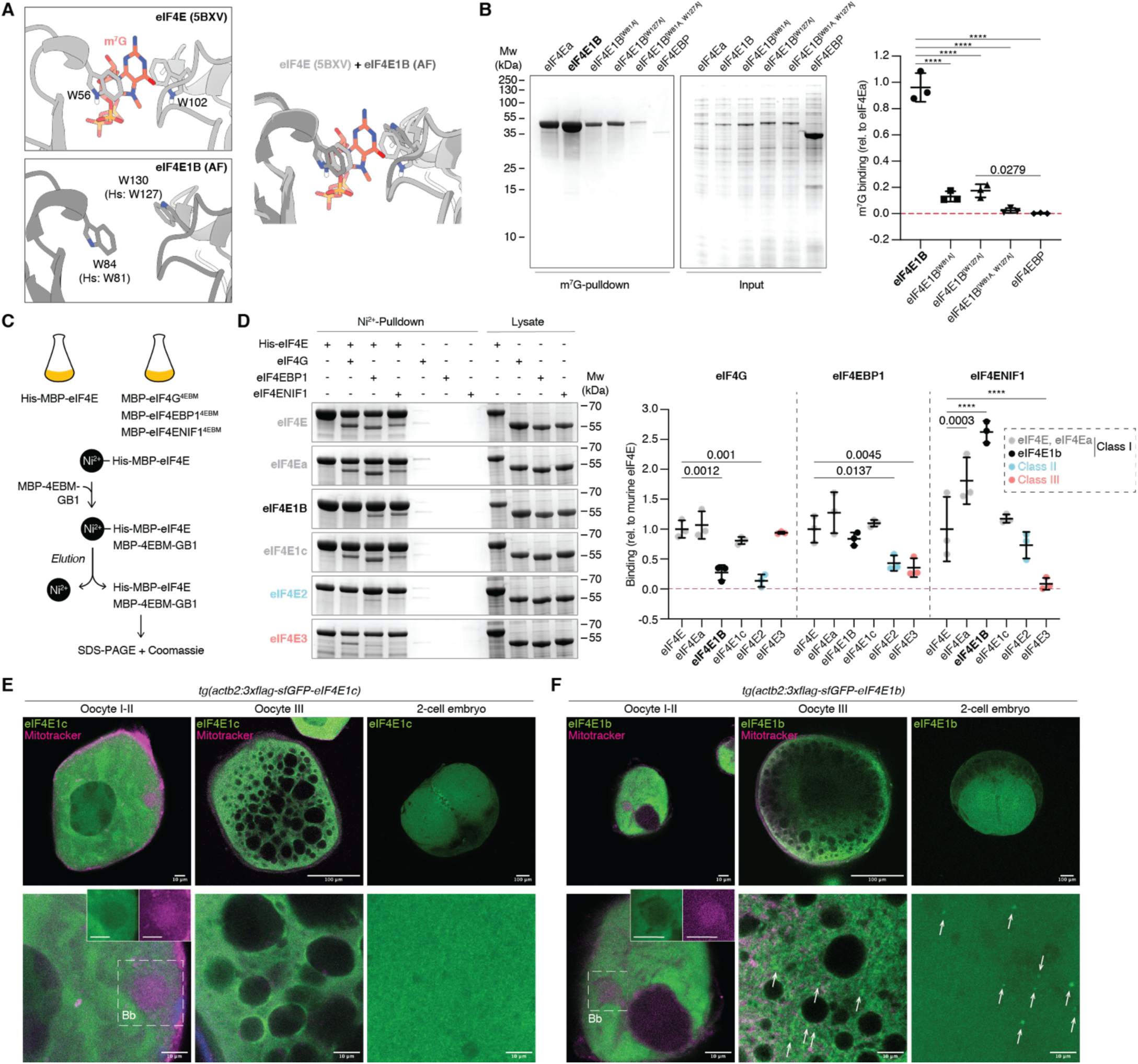
eIF4E1b interacts with the mRNA cap, eIF4EBP1 and eIF4ENIF1 and localizes to cytoplasmic foci in oocytes and embryos. **A** eIF4E interaction with the mRNA cap (top, PDB-5BXV, Sekiyama *et al*, 2015) is mediated by two tryptophans that are conserved in eIF4E1B (bottom, AlphaFold (AF) prediction of mouse eIF4E1B). The superimposition of both structures is shown on the right, with eIF4E in light gray, eIF4E1b in dark gray, and m^7^G in red. **B** (Left) Coomassie-stained gels from immunoprecipitation assays with *E. coli* lysates containing zebrafish eIF4Ea and human eIF4E1B (wild type and tryptophan mutants, see *A*) using m^7^G-coated beads. The eIF4E-binding motif of human eIF4EBP1 is used as negative control. Quantification of eIF4E binding to m^7^G (relative to eIF4Ea) is shown on the right. **C** Scheme of in vitro pulldown assays. Lysates of *E. coli* cells expressing His- and MBP-tagged eIF4Es were incubated with Ni^2+^ beads. Lysates containing MBP-tagged eIF4E-binding motifs (4EBM) of human eIF4G, eIF4EBP1 or eIF4ENIF1 (see **Fig EV4C-E**) were added to the beads. After elution, binding of eIF4G, eIF4EBP1 and eIF4ENIF1 to eIF4E was assessed by SDS-PAGE and Coomassie staining. **D** (Left) Coomassie-stained gels of pulldowns with mouse eIF4E, eIF4Es from zebrafish (eIF4Ea, eIF4E2 and eIF4E3), and mouse eIF4E1B with 4EBMs from eIF4G, eIF4EBP1 and eIF4ENIF1. Quantifications are shown on the right. **E-F** Confocal microscopy images of fixed transgenic zebrafish oocytes and embryos expressing 3xflag-sfGFP-eIF4E1c (*E*) and 3xflag-sfGFP-eIF4E1b (*F*). Mitochondria were stained with Mitotracker (in magenta). Images at two different magnifications are shown at the top and bottom (scale bars correspond to 100 and 10 μm, respectively). The Balbiany body (Bb) is indicated by a dashed box; individual channels are shown in boxes for the Bb. Cytoplasmic foci are depicted with arrows. Data information: In *B* and *D*, n = 3 independent experiments. Significance was determined using two-way ANOVA (*B*) or one-way ANOVA (*D*) followed by Tukey’s (*B*) or Dunnett’s (*D*) multiple comparisons test (****: p-value < 0.0001).

The interaction of eIF4E with eIF4ENIF1 has been reported to trigger the localization of eIF4E to P-bodies (Ferraiuolo *et al*, 2005). Since eIF4E1b showed the highest affinity for eIF4ENIF1 among all eIF4Es tested, we investigated the subcellular localization of eIF4E1b. To this end, we generated transgenic zebrafish lines expressing GFP-tagged versions of eIF4Ea, eIF4E1c and eIF4E1b under the control of the ubiquitously expressed *β-actin* promoter. GFP-tagged eIF4Es localized to the cytoplasm of zebrafish oocytes and were excluded from the Balbiani body (**Fig 2E and F, Fig EV5A**), a membraneless organelle composed of mitochondria, endoplasmic reticulum and RNA that is important for germline determination in zebrafish (Jamieson-Lucy & Mullins, 2019a). While eIF4Ea and eIF4E1c showed a diffuse cytosolic signal at all stages of oogenesis (**Fig 2E, Fig EV5A**), eIF4E1b formed cytoplasmic puncta in stage III oocytes (**Fig 2F**), similar to the localization observed for the P-body component DDX6 in mouse oocytes (Flemr *et al*, 2010). In activated eggs and embryos, eIF4E1b localized to the cytosol and to cytoplasmic granules colocalizing with P-body markers such as Dcp2, Ddx6 and Ybx1 (**Fig 2F, Fig EV5B-D**). These data suggest that eIF4E1b localizes to P-bodies during embryogenesis, in line with the interaction observed between eIF4E1b and eIF4ENIF1 in vitro.

Although eIF4E1b shares ∼60% amino acid identity with other class I eIF4Es, our in vitro and in vivo data suggest that eIF4E1b does not function as a canonical eIF4E. We combined sequence alignments with available structural data and AlphaFold (AF) predictions (Mirdita *et al*, 2022; Evans *et al*, 2022) of mouse eIF4E paralogs to investigate the contribution of specific amino acids in determining eIF4E1B interactions. AF predicted an eIF4E1B structure very similar to eIF4E (**Fig EV6A**). The eIF4E-binding motifs of eIF4G, eIF4EBP and eIF4ENIF1 have been reported to bind to the so-called dorsal and lateral surfaces of eIF4E (**Fig 3A**) (Igreja *et al*, 2014; Sekiyama *et al*, 2015; Peter *et al*, 2015; Grüner *et al*, 2018). Mutation of the conserved Trp84 located on the dorsal surface of mouse eIF4E1B (**Fig 3B**) abolished the interaction with eIF4EBP1 and eIF4ENIF1 (**Fig 3D, Fig EV6B**), indicating a similar binding mode of eIF4E-binding motifs to the dorsal surfaces of eIF4E and eIF4E1B. Mutation of residues located on the lateral surface of mouse eIF4E1B (**Fig 3B**) abolished eIF4EBP1 binding, whereas eIF4ENIF1 binding was only reduced (**Fig 3D, Fig EV6B**), suggesting that additional residues in eIF4E1B stabilize the interaction with eIF4ENIF1. To investigate this further, we generated eIF4E1B-eIF4E chimeras (**Fig EV2A**). Exchanging the N-terminal half of eIF4E1B (eIF4E-eIF4E1B) restored its ability to interact with eIF4G and decreased its affinity for eIF4ENIF1 (**Fig 3D, Fig EV6B**), suggesting that the N-terminal half of eIF4E1B is important for defining eIF4E1B interactions. Based on AF predictions, we hypothesized that Lys108 of mouse eIF4E1B binds to eIF4ENIF1 (**Fig 3E**), reminiscent of the interaction observed between Drosophila eIF4E and the fly ortholog of eIF4ENIF1, 4E-T (**Fig EV6C**) (Peter *et al*, 2015). In eIF4Es, an Asn is predicted to form an intramolecular amide bond with the residue equivalent to Lys108 in mouse eIF4E, Gln80, thereby preventing the interaction of this residue with eIF4ENIF1 (**Fig 3E**). In line with this hypothesis, mutation of Lys108 to an opposite charge (Glu) or of Lys112 to an Asn (as in eIF4Es) reduced the affinity of eIF4E1B for eIF4ENIF1 (**Fig 3D, Fig EV6B**). In addition to Lys112, we identified three other residues within the N-terminal half of eIF4E1Bs that are conserved in eIF4Es but differ in eIF4E1Bs (**Fig 3C, Fig EV2A**). However, mutation of all four residues to their equivalent amino acids in eIF4Es did not affect eIF4G binding, whereas it decreased the affinity for eIF4ENIF1 to a similar extent as mutation of Lys112 alone (**Fig 3D, Fig EV6B**). Taken together, our in vitro data show that eIF4E1Bs contain specific residues that interact with eIF4ENIF1 and increase its affinity for this protein.

**Figure 3.**
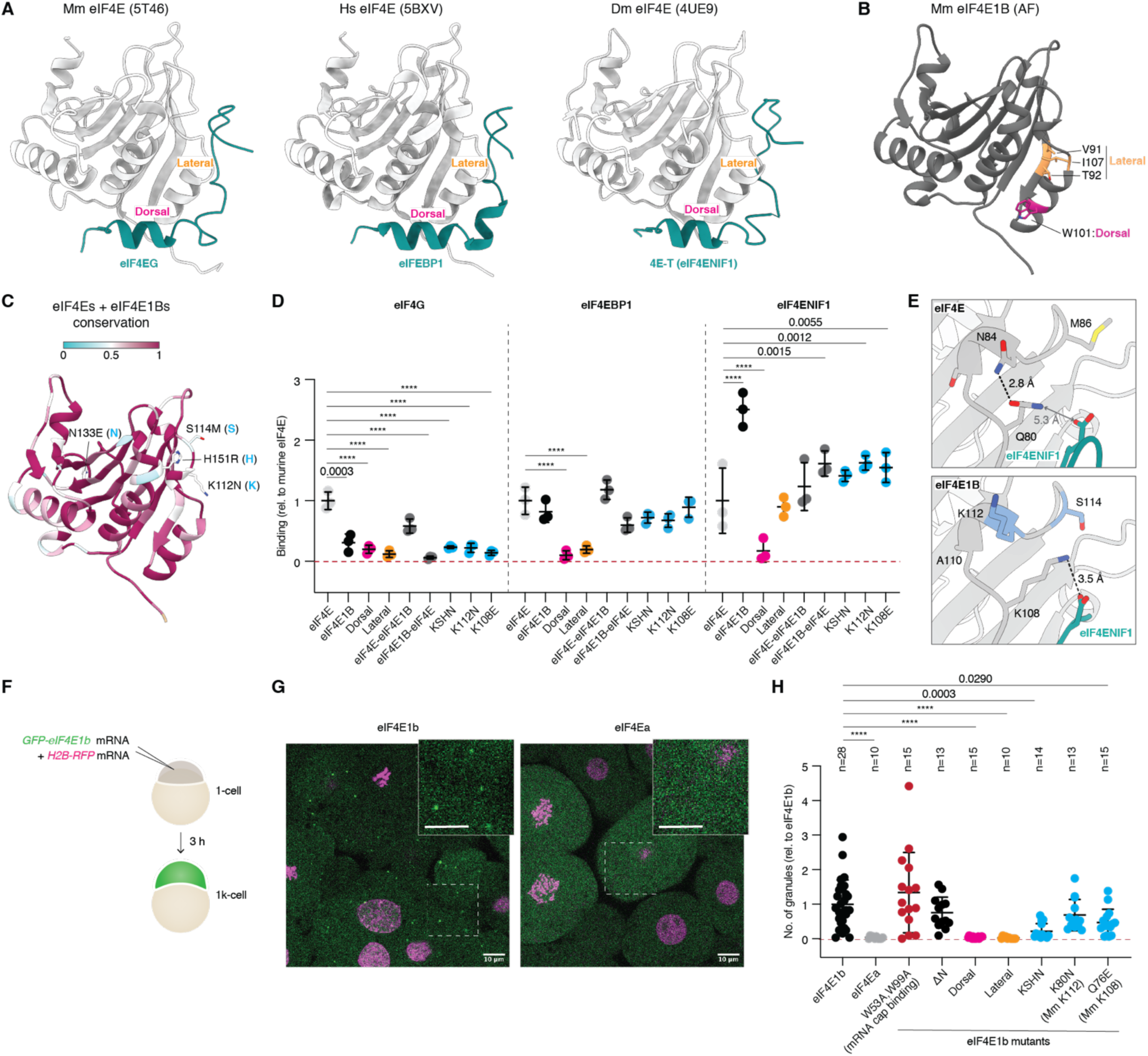
Specific residues in eIF4E1b increase its affinity for eIF4ENIF1 and promote its localization to P-bodies in the embryo. **A** Structures of eIF4E proteins (in gray) bound to the eIF4E-binding motifs (in blue) of eIF4G (PDB-5T46; Grüner *et al*, 2016), eIF4EBP1 (PDB-5BXV; Sekiyama *et al*, 2015), and 4E-T/eIF4ENIF1 (PDB-4UE9; Peter *et al*, 2015). **B** AlphaFold (AF)-predicted structure of mouse eIF4E1B. Residues located at the dorsal and lateral surfaces are highlighted in magenta and orange, respectively. **C** The AF structure of mouse eIF4E1B is colored based on amino acid conservation among vertebrate eIF4E and eIF4E1B proteins (sequences in **Fig EV2A**). Residues in the N-terminal half that differ in eIF4E1Bs but are conserved in eIF4Es are indicated (first residue: eIF4E1b; second residue: eIF4E). **D** Quantification of eIF4G, eIF4EBP1 and eIF4ENIF1 binding to mouse eIF4E1B wild-type and mutant proteins in pulldowns with *E. coli* lysates (see **Fig EV6B**; n = 3 independent experiments), compared to mouse eIF4E. eIF4E and eIF4E1B data are also plotted in **Fig 2D**. KSHN refers to the residues highlighted in *C*. **E** AF-predicted structures of mouse eIF4E (top) or eIF4E1B (bottom) in complex with the eIF4E-binding motif of human eIF4ENIF1. Distances are indicated; interactions are depicted with dashed lines. **F** Assay to test the contribution of specific amino acids in determining the subcellular localization of eIF4E1b in zebrafish embryos. mRNAs were co-injected into 1-cell embryos; embryos were imaged after 3 hours. **G** Representative confocal microscopy pictures of live embryos transiently expressing GFP-tagged eIF4E1b or eIF4Ea from mRNAs. Regions delimited by dashed boxes are shown with a higher magnification (scale bars = 10 μm). **H** Number of eIF4E-positive granules relative to wild-type eIF4E1b (n = embryos). Data information: In *D* and *H*, significance was determined with two-way (*D*) or one-way (*H*) ANOVA followed by Dunnett’s test (****: p-value < 0.0001).

eIF4ENIF1 has been reported to target eIF4E to P-bodies in human cells (Ferraiuolo *et al*, 2005). Since eIF4E1B interacts with eIF4ENIF1 in vitro and localizes to P-bodies in zebrafish embryos (in contrast to the cytosolic and nuclear localization of other class I eIF4Es with lower affinities for eIF4ENIF1; **Fig 2D and E, Fig EV5B**), eIF4ENIF1 may also determine the subcellular localization of eIF4E1b in zebrafish. Moreover, binding to the mRNA cap or to other proteins may also influence the assembly of eIF4E1b into P-bodies. To test these ideas, we expressed GFP-tagged wild-type or mutant versions of zebrafish eIF4Es via mRNA injections into 1-cell embryos and imaged their localization at 3 hpf (**Fig 3F and G**).

Localization of zebrafish eIF4E1b to P-bodies was not affected by mutation of Trp53 and Trp99, which are required for mRNA cap binding in vitro (**Fig 2A and B**), or by deletion of its unstructured N-terminus, which is the region most distinct from eIF4Es (**Figs 1A and 3H, Fig EV6D**). In contrast, mutation of residues located on the dorsal or lateral surface of eIF4E1b that are important for the interaction with eIF4EBP and eIF4ENIF1 abolished its localization to P-bodies (**Fig 3H, Fig EV6D**). Notably, mutation of residues identified in our in vitro experiments as important for eIF4ENIF1 but not for eIF4EBP1 binding (see **Fig 3D**) was sufficient to reduce the number of eIF4E1b foci in the embryo (**Fig 3H, Fig EV6D**). Taken together, these data suggest that eIF4E1b does not have an intrinsic ability to localize to P-bodies, but it does so through interaction with eIF4ENIF1.

To gain insights into the eIF4E1b interactome, we performed immunoprecipitation (IP) followed by mass spectrometry experiments using anti-GFP beads and lysates from either wild-type or transgenic zebrafish oocytes and embryos expressing GFP-tagged eIF4E1b. We used transgenic zebrafish expressing GFP-tagged eIF4E1c as a control because eIF4E1c is abundant during early embryogenesis (**Fig 1D**) and has a similar affinity for eIF4G, eIF4EBP1, and eIF4ENIF1 as mammalian eIF4E in vitro (**Fig 2D**). In oocytes and 8-cell embryos, eIF4E1b interacted with eIF4EBPs and P-body components, including eIF4ENIF1, Zar1l/Zar2, Lsm14/Rap55 proteins and Ddx6 (**Fig 4A and B, Fig EV7A**). Notably, all P-body components interacted with eIF4E1b only in the presence of RNA except eIF4ENIF1 (**Fig EV7C**), suggesting that eIF4ENIF1 is the only protein that directly binds to eIF4E1b. At 3 hpf, during zygotic genome activation (ZGA), eIF4E1b no longer interacts with Zar1l, whose mRNA and protein levels decline after 1 hpf (**Fig 4C, Fig EV7D and E**). In line with our in vitro experiments, eIF4E1c interacted with eIF4Gs, eIF4EBPs and eIF4ENIF1 in vivo (**Fig 4D-F, Fig EV7B**). Moreover, translational factors belonging to the eIF3 complex were specifically enriched in the eIF4E1c IP at 3 hpf (**Fig 4F**), consistent with the increase in translation observed during the first hours of zebrafish embryogenesis (Leesch *et al*, 2023). These results suggest that eIF4E1c functions as a canonical eIF4E in vivo, promoting translation initiation, whereas eIF4E1b plays a role in P-bodies. While mRNA decapping components (e.g. Dcp1, Dcp2, and Edc4) localize to P-bodies (Greber *et al*, 2016) (**Fig EV5D**), it is of noteworthy that none of these proteins were detected in eIF4E1b IPs, despite their expression in zebrafish oocytes and embryos (**Fig EV7F-H**). This suggests that eIF4E1b interferes with the binding of the decapping machinery to the mRNA cap, as reported for canonical eIF4E in human cells and yeast (Schwartz & Parker, 2000; Räsch *et al*, 2020), consistent with the similar binding mode of eIF4E and eIF4E1B to the mRNA cap (**Fig 2A and B**).

**Figure 4.**
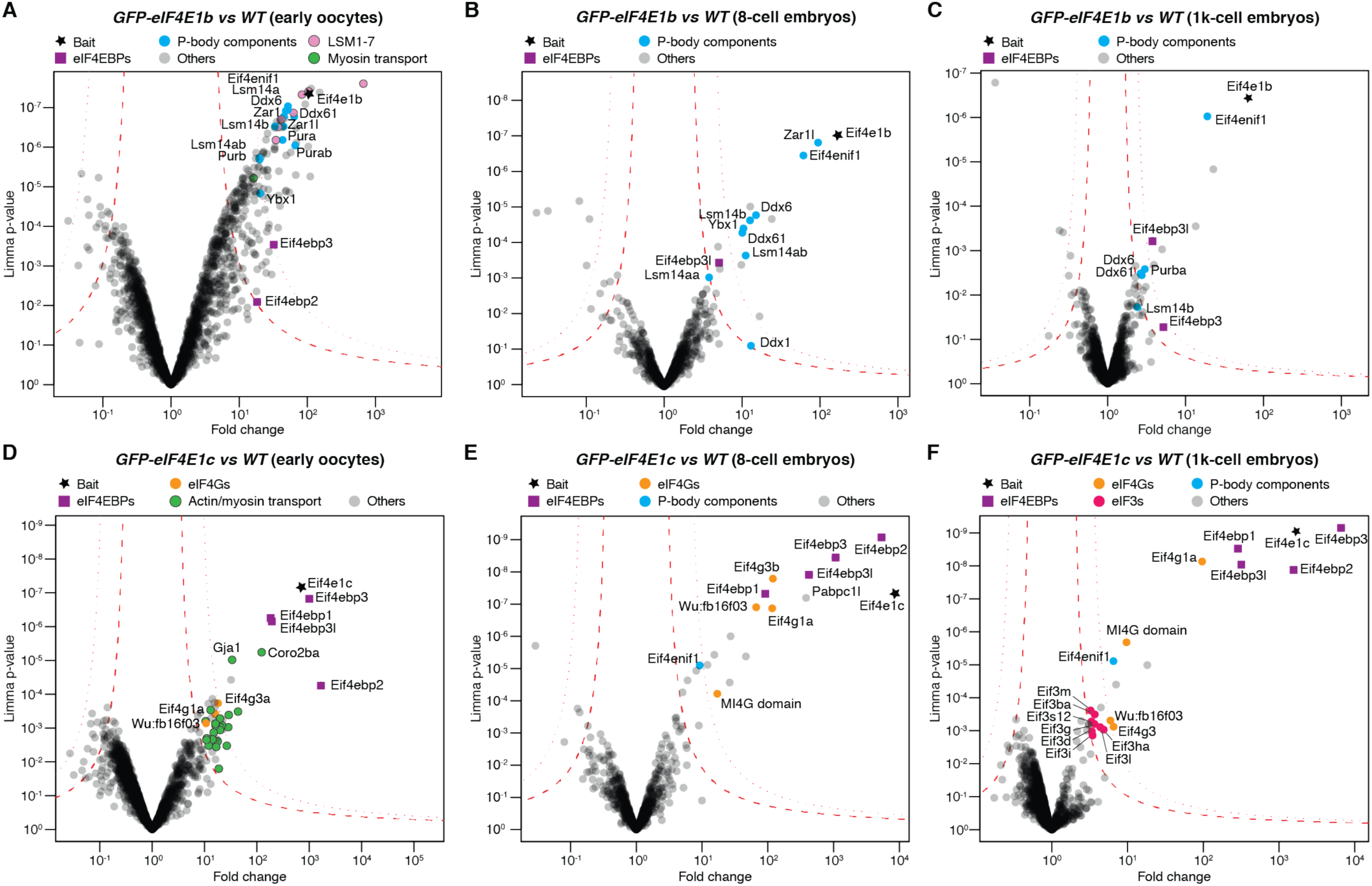
eIF4E1b interacts with P-body components in zebrafish oocytes and embryos. Volcano plots of immunoprecipitation followed by mass spectrometry (IP-MS) experiments with anti-GFP beads using lysates from wild-type (*WT*) and transgenic oocytes and embryos expressing GFP-tagged eIF4E1b (*A-C*) or eIF4E1c (*D-F*). MS experiments were performed from early oocytes (oogonia and stages I-II; *A* and *D*) and embryos (*B, C*, *E*, and *F*). Permutation-based false discovery rates (FDRs) are displayed as dotted (FDR < 0.01) and dashed (FDR < 0.05) lines (n = 3 biological replicates).

To identify which mRNAs are bound by eIF4E1b, we performed RNA immunoprecipitation (RIP) followed by sequencing in transgenic zebrafish 8-cell embryos expressing GFP-tagged eIF4E1b or eIF4E1c. Differential expression analysis revealed 1206 and 1414 genes as specifically enriched in eIF4E1b and eIF4E1c RIPs, respectively (p-value < 0.005) (**Fig 5A, Fig EV8A and B**). Overexpression of eIF4E1b and eIF4E1c did not result in major changes in the transcriptome, with 1.6% (363 transcripts) and 2.2% (500 transcripts) being differentially enriched in lysates from 8-cell embryos overexpressing eIF4E1b and eIF4E1c, respectively (**Fig EV8C-F**). A large fraction of the transcripts upregulated in eIF4E1b or eIF4E1c overexpressing embryos was also enriched in the eIF4E1b or eIF4E1c RIP, respectively (**Fig EV8G**), suggesting a role for both eIF4Es in stabilizing the mRNAs to which they bind. Genes enriched in the eIF4E1b RIP had GO-terms related to chromatin regulation, whereas genes enriched in the eIF4E1c RIP belonged to GO-terms associated with mRNA processing and export (**Fig 5B**). Consistent with our data suggesting a role for eIF4E1b in mRNA storage and for eIF4E1c in translation initiation, mRNAs involved in chromatin organization and remodeling have been reported to be deadenylated and stable during the oocyte-to-egg transition in mouse (Lee *et al*, 2023), whereas mRNAs involved in mRNA processing have been reported to be translated upon fertilization in sea urchin (Chassé *et al*, 2018). To investigate the translational status of the mRNAs bound by eIF4E1b and eIF4E1c in zebrafish embryos, we compared our RIP-seq data with publicly available datasets of mRNAs expressed during zebrafish embryonic development (Cabrera-Quio *et al*, 2021; Chang *et al*, 2018; Subtelny *et al*, 2014). Of note, mRNAs enriched in eIF4E1b RIP were overall less abundant than mRNAs enriched in eIF4E1c RIP in RNA-seq datasets obtained with polyA-selected and rRNA depletion (RiboMinus) protocols (Cabrera-Quio *et al*, 2021) (**Fig 5C**). Analysis of polyA tail length (Chang *et al*, 2018) revealed that most transcripts enriched in the eIF4E1b RIP had short polyA tails during the first 4 hpf, in stark contrast to the long polyA tails observed for mRNAs enriched in the eIF4E1c RIP (**Fig 5D**). Consistent with mRNAs with long polyA tails being specifically enriched in eIF4E1c RIP, our IP-MS experiments showed a >300-fold enrichment of the cytoplasmic polyA-binding protein PABPC1L in eIF4E1c IPs from 8-cell embryos, whereas it was only < 3.5-fold enriched in eIF4E1b IPs. Following the general trend of maternal transcripts, the polyA tails of mRNAs enriched in the eIF4E1b RIP also increased in length during embryogenesis (**Fig 5C and D**). In line with the strong correlation observed between polyA tail length and translation during the first hours of embryogenesis (Subtelny *et al*, 2014), as well as with our data showing that eIF4E1b does not interact with eIF4G, mRNAs enriched in the eIF4E1b RIP were translationally repressed according to previously published ribosome profiling data (**Fig 5E**) (Subtelny *et al*, 2014). In contrast, mRNAs enriched in the eIF4E1c RIP showed a high translational efficiency (TE) (**Fig 5E**) (Subtelny *et al*, 2014). Taken together, our RIP-seq data and analyses suggest that eIF4E1b binds to translationally repressed mRNAs with short polyA tails. The median polyA tail length of mRNAs enriched in the eIF4E1b RIP is 12 nucleotides in the 1-cell embryo, which is 25% shorter than the median polyA tail of all maternal mRNAs at the same developmental stage (Chang *et al*, 2018). In somatic cells, shortening of polyA tails to 10-12 nucleotides promotes mRNA decapping and degradation (Passmore & Coller, 2022). However, maternal mRNAs with polyA tails shorter than 12 nucleotides are stable during the first hours of embryogenesis (**Fig EV8H**) (Bhat *et al*, 2023). Our data suggest that eIF4E1b binds to a subset of maternal mRNAs with short polyA tails to promote their storage and translational repression, thus contributing to the uncoupling of polyA tail shortening and mRNA decay observed during early embryogenesis (**Fig 5F**).

**Figure 5.**
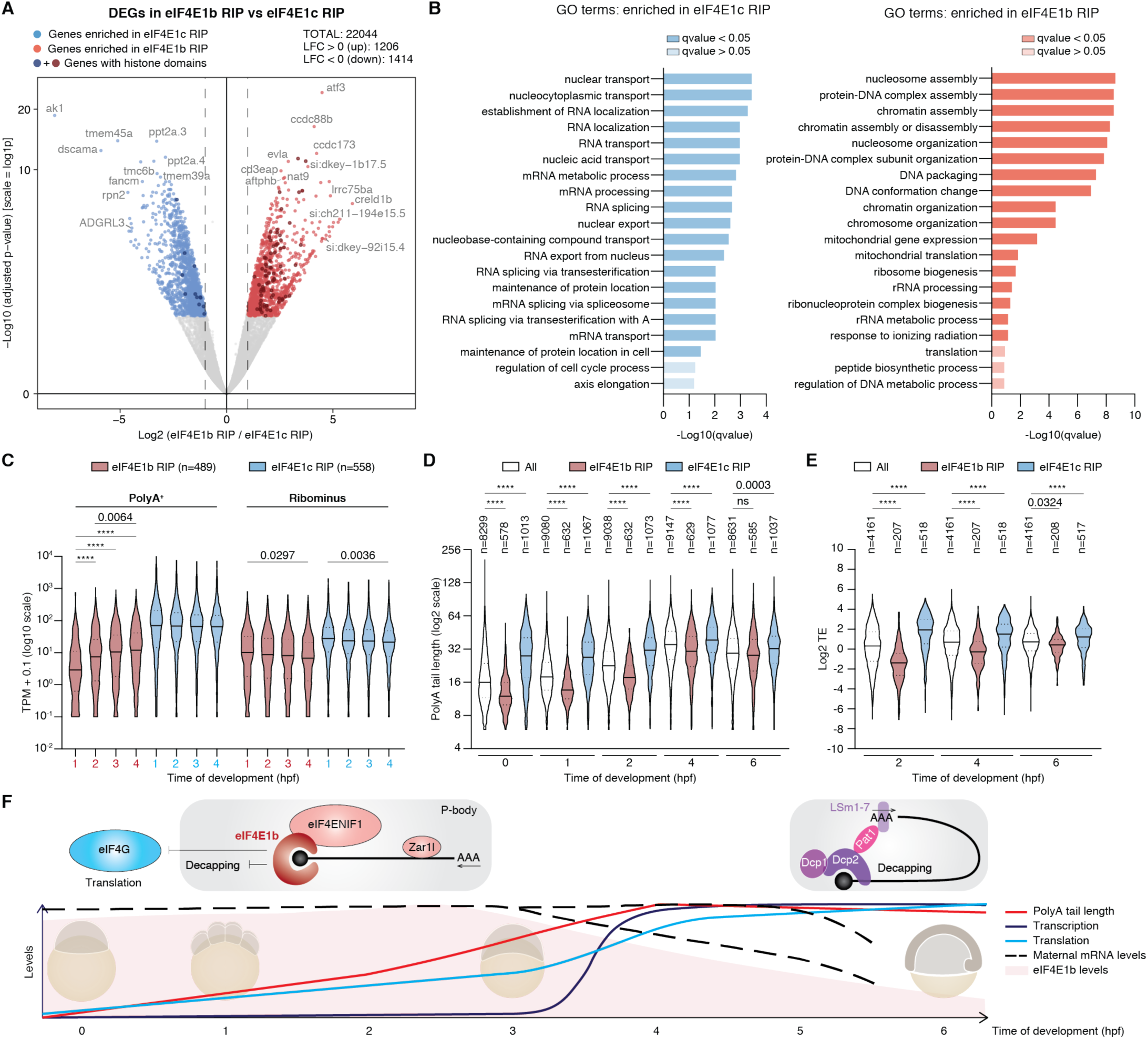
eIF4E1b binds to translationally repressed mRNAs involved in chromatin regulation. **A** Differential expression gene (DEG) analysis of mRNAs enriched in RNA immunoprecipitation experiments (RIPs) with eIF4E1b (in red; n = 3 biological replicates) versus those enriched in eIF4E1c RIPs (in blue; n = 2 biological replicates) in 8-cell transgenic embryos expressing sfGFP-tagged eIF4E1b or eIF4E1c (p-value < 0.005). Genes encoding proteins with histone and histone-like domains are highlighted in a darker color. **B** Gene ontology (GO) analysis of mRNAs that are specifically bound to eIF4E1c (left, in blue) or to eIF4E1b (right, in red). **C** Expression levels (in transcript per million, TPM) of mRNAs enriched in eIF4E1b and eIF4E1c RIPs during zebrafish embryogenesis according to published RNA-seq datasets obtained with polyA-selection (PolyA^+^) and rRNA depletion (Ribominus) protocols (Cabrera-Quio *et al*, 2021). **D** PolyA tail length of mRNAs enriched in RIPs with eIF4E1b or eIF4E1c during zebrafish embryonic development based on published TAIL-seq data (Chang *et al*, 2018) (ns: non-significant). **E** Translational efficiency (TE) of mRNAs enriched in eIF4E1b and eIF4E1c RIPs during zebrafish embryogenesis based on published TE data (Subtelny *et al*, 2014). **F** Cartoon depicting eIF4E1b function during early zebrafish embryogenesis. Maternal mRNAs with short polyA tails are translationally repressed in the egg (bottom; Subtelny *et al*, 2014). eIF4E1b interacts with eIF4ENIF1 and localizes to P-bodies, where it binds to deadenylated mRNAs that are also bound by Zar1l (top left). Binding of eIF4E1b to the mRNA cap interferes with decapping and with eIF4F-dependent translation. Maternal mRNA clearance (according to Bhat *et al*, 2023) coincides with a decrease in eIF4E1b levels, which may then allow the decapping and degradation of mRNAs with short polyA tails (top right). Data information: In *C*-*E*, significance was assessed with Kruskal-Wallis followed by Dunn’s test (****: p-value < 0.0001). For *C*, significance was calculated for eIF4E1b RIP or eIF4E1c RIP in PolyA^+^ or Ribominus data (if not stated, p-value > 0.005). For *D* and *E*, statistics are only shown for comparisons within the same developmental stage.

## Discussion

Our in vivo and in vitro data reveal that eIF4E1b is a non-canonical eIF4E that plays a critical role in female germline development in zebrafish, in line with a recent study in mice showing that eIF4E1B is essential for female fertility (Guo *et al*, 2023). While eIF4E1b binds to the mRNA cap like other class I eIF4Es, it does not interact with the translation initiation factor eIF4G and is therefore not involved in initiating eIF4F-dependent translation. In yeast, eIF4G interacts with Dcp1 and may thus recruit the decapping machinery to mRNAs (Vilela *et al*, 2000). If this interaction also exists in vertebrates, eIF4E1b may inhibit decapping not only by blocking access for Dcp2 to the mRNA cap, similarly to other eIF4Es (Schwartz & Parker, 2000), but also by interfering with eIF4G binding to mRNAs. Moreover, our data show that eIF4E1b has a strong affinity for eIF4ENIF1, a P-body component that targets eIF4E1b to P-bodies in the embryo. Binding of eIF4ENIF1 to eIF4Es has been shown to stabilize deadenylated mRNAs in human cells (Räsch *et al*, 2020). Mutations in *eIF4ENIF1* also cause infertility in women (Kasippillai *et al*, 2013; Zhao *et al*, 2019; Shang *et al*, 2022), in agreement with both eIF4E1B and eIF4ENIF1 working together to promote maternal mRNA dormancy.

Previous studies suggest that mRNAs localizing to P-bodies are translationally repressed (Brengues *et al*, 2005; Halstead *et al*, 2015; Hubstenberger *et al*, 2017). Consistent with the localization of eIF4E1b to P-bodies during early zebrafish embryogenesis, mRNAs enriched in eIF4E1b RIP from 8-cell embryos have low translational efficiency. Our in vitro and in vivo data further demonstrate that eIF4E1Bs do not bind to eIF4G and cannot be involved in promoting eIF4F-dependent translation, as recently proposed for mouse eIF4E1B (Guo *et al*, 2023). Instead, we find that eIF4E1Bs directly interact with the eIF4E-binding motifs of eIF4EBP1 and eIF4E1NIF1, contradicting previous studies that reported no binding of eIF4E1Bs to these sites (Robalino *et al*, 2004; Minshall *et al*, 2007). We also observed a strong binding of eIF4E1B to the mRNA cap (in contrast to Robalino *et al*, 2004; Minshall *et al*, 2007; Kubacka *et al*, 2015), which may be important for its function in inhibiting mRNA decapping.

Interestingly, histone mRNAs were particularly enriched in the eIF4E1b RIP (**Fig 5A**; 77 genes enriched in the eIF4E1b RIP versus 12 in the eIF4E1c RIP). Consistent with this finding, we identified the stem loop binding proteins SLBP and SLBP2, which bind to the 3’ UTR of histone mRNAs, enriched in eIF4E1b IPs from oocytes and 8-cell stage embryos, respectively. While maternal histones are deposited in lipid droplets in Drosophila (Li *et al*, 2012), similar droplets have so far not been reported in vertebrate embryos. In somatic cells, histone mRNA synthesis and degradation are coupled to the G1/S phase of the cell cycle (Graves *et al*, 1987; Armstrong & Spencer, 2021). The absence of transcription during early embryogenesis raises the exciting possibility that histone mRNAs, which are highly abundant during the first hours of embryogenesis (**Fig EV8I**), are stored in the egg for later translation in the embryo. Since histone mRNAs lack a polyA tail, they cannot be repressed by deadenylation like other maternal mRNAs. eIF4E1b may therefore play a critical role in repressing translation of histone mRNAs, thereby preventing the toxicity caused by an excess of histones in cells (Singh *et al*, 2010). Furthermore, histone mRNA decay is coupled to its translation (Stimac *et al*, 1983; Tuck *et al*, 2020), supporting an additional role for eIF4E1b in stabilizing maternal histone mRNAs.

One key open question is what determines the selectivity of eIF4E1b for certain mRNAs. According to TAIL-seq data (Chang *et al*, 2018), more than 2000 genes have mRNAs with a median polyA tail length of less than 12 nucleotides in the zebrafish egg. However, only a fraction of these genes (276) correspond to transcripts enriched in eIF4E1b RIP, suggesting that (1) eIF4E1b is not recruited to every mRNA that contains a short polyA tail and (2) mRNAs with short polyA tails may also be stabilized by other mechanisms. What determines the specificity of eIF4E1b for certain mRNAs and how mRNAs are released from eIF4E1b to enable their translation during embryogenesis remains to be discovered. While answering these questions will require future work, the conservation of eIF4E1b in vertebrates and its expression in the human brain (Human Protein Atlas version 22.0) suggest that our findings from the zebrafish oocyte-to-embryo transition may have direct relevance in other systems and cellular contexts.

## Materials and Methods

### Phylogenetic analysis

Sequences were collected from the UniProt or NCBI protein databases with NCBI blast+ (Camacho *et al*, 2009) and aligned with mafft (v7.505, -linsi method) (Katoh & Toh, 2008). For phylogenetic analysis, a maximum likelihood tree was inferred with iqtree2 v.2.2.0 (Minh *et al*, 2020), with standard model selection using ModelFinder (Kalyaanamoorthy *et al*, 2017) and ultrafast bootstrap (UFBoot2) support values (Hoang *et al*, 2018). The tree was visualized in iTOL v6 (Letunic & Bork, 2021).

### Tandem mass tag mass spectrometry (TMT-MS)

#### Sample preparation

Embryos were manually dechorionated and deyolked in Danieau’s buffer, containing 58 mM NaCl, 0.7 mM KCl, 0.4 mM MgSO4, 0.6 mM Ca(NO_3_)_2_ and 5 mM HEPES pH 7.2, and snap frozen in liquid nitrogen. 64 embryos per stage from 16-19 females were shacked at 800 rpm for 5 min at 95°C in 4% SDS, 0.1 MDTT, 0.1 M Tris-HCl pH 7.5, and 300 pmol of 7 recombinantly expressed proteins (dCas9, SpoIVB deltaN34 from *G. stearothermophilus*, ClpC NTD from *B. subtilis*, lambda exonuclease, YwlE from *G. stearothermophilus*, McsA from *S. aureus* and SpoIVFAGs.deltaN137_L234M; Molecular Biology Service, IMP). Lysates were sonicated and the SDS was removed using filter-aided sample preparation (FASP) (Wiśniewski *et al*, 2009). Proteins were digested using Trypsin Gold (Promega) in 100 mM HEPES pH 7.6 at 37°C overnight. Samples were acidified with 10% TFA and purified using a Sep-Pak C18 Vac cartridge (Waters). Peptides were dissolved in 100 mM HEPES pH 7.6. Equal volumes of lysates were incubated with TMT10plex reagents for 2 h. After adding 5% hydroxilamin, samples were purified using a Sep-Pak C18 Vac cartridge (Waters), dried, and dissolved in SCX buffer A (5 mM phosphate buffer pH 2.7, 15% ACN).

#### LC-MS

MS analysis was performed on an UltiMate 3000 RSLCnano system (ThermoFisher) coupled to a Q Exactive HF-X mass spectrometer (ThermoFisher) with a Proxeon nanospray ion source (ThermoFisher) as described in Leesch *et al*, 2023. The mass spectrometer was operated in data-dependent mode, with a full scan (m/z range 380–1650, resolution of 120000, target value 3E6), followed by MS/MS scans of the 10 most abundant ions. MS/MS spectra were acquired using a normalized collision energy of 35, isolation width of 0.7 m/z, resolution of 45000, a target value of 1E5 and maximum fill time of 250 ms. For the detection of the TMT reporter ions, a fixed first mass of 110 m/z was set. Precursor ions selected for fragmentation (excluding ions with charge state 1, 7, 8, >8) were put on an exclusion list for 30 sec. The minimum AGC target was set to 1E4 and intensity threshold was calculated to be 4E4.

#### TMT-MS data analysis

Peptide spectra were identified with Proteome Discoverer (v2.3.0.523, ThermoFisher) and searched against a custom-made protein database covering GRCz10, GRCz11, RefSeq, UniProt and PDB identifiers (58522 sequences; 34078760residues), using MS Amanda (v2.0.0.14114) (Dorfer *et al*, 2014) with the following parameters: iodoacetamide derivative on cysteine was set as a fixed modification, whereas oxidation on Met, deamidation on Asn and Gln, phosphorylation on Ser, Thr and Tyr and TMT10plex on Lys and peptide N-termini were set as variable modifications. Monoisotopic masses were searched within unrestricted protein masses for tryptic enzymatic specificity. The peptide mass tolerance was set to ± 5 parts per million (ppm) and the fragment mass tolerance to ± 15 ppm. The maximal number of missed cleavages was set to 2. Results were filtered to 1% false discovery rate (FDR) on protein level using the Percolator algorithm integrated in Proteome Discoverer (Käll *et al*, 2007). Peptides were quantified based on TMT reporter ion intensities using the Reporter Ion Quantifier Node in Proteome Discoverer.

### Zebrafish husbandry

Zebrafish (*Danio rerio*) were raised at 28°C with a 14/10 hour of light/dark cycle. Wild-type TLAB fish correspond to fish obtained by crossing AB with the natural variant TL (TupfelLongfin). All fish experiments were conducted according to Austrian and European guidelines for animal research and approved by local Austrian authorities (protocols for work GZ342445/2016/12 and MA 58-221180-2021-16).

### Zebrafish knockout and transgenic lines

Zebrafish *eif4e1b* knockout fish were generated by CRISPR-Cas9 mediated mutagenesis. Guide-RNAs (gRNAs) targeting the third exon of *eIF4E1B* (5’-CACCAAATTCGACACGGTCGAGG-3’ and 5’-GACACGGTCGAGGACTTCTGGGG-3’) were injected together with recombinant Cas9 protein (Molecular Biology Service, IMP) into 1-cell zebrafish embryos. To identify fish carrying mutations in the germ line (i.e. founders), adult fish were crossed to wild type; embryos were genotyped by PCR using 5’-AGATGGGGACTTTGGTTCTACA-3’ and 5’-TGCTCTACTCCACCTTTCACAA-3’ primers. Embryos showing a size difference in PCR amplicons were raised to adulthood and heterozygous fish were identified by PCR-based genotyping of fin clips. Homozygous fish and their wild-type siblings were generated by crossing heterozygous fish. Sex was determined at 4-5 months post fertilization based on the sexual dimorphism characteristic of zebrafish. Pictures of anesthetized fish (in MESAB) were taken on a ZEISS Stemi 508 stereo microscope with camera (2x magnification, FlyCapture2 software). Fertilization rates were counted at 3 hpf; unfertilized eggs were removed, and embryo survival was counted at 1 dpf. Graphs and statistical analyses were performed using GraphPad Prism 8.0.2.

To generate transgenic lines, the coding sequences of full length eIF4Ea, eIF4E1b and eIF4E1c were PCR-amplified from zebrafish cDNA and cloned by Gibson assembly into a vector containing Tol2-integrations sites in between the zebrafish *actb2* promoter, *actb2* 5’ UTR, an N-terminal *3xflag-sfGFP* tag and the SV40 late polyadenylation signal (*SVLPA*). 15 pg of each plasmid was co-injected with 35 pg of *Tol2* mRNA into 1-cell embryos. Adult fish were crossed with wild-type fish, and GFP-positive embryos were raised to adulthood. Rescue lines were obtained by injecting the *3xflag-sfGFP-eIF4E1B* vector into *eif4e1b* heterozygous embryos. Homozygous fish and wild-type fish were obtained by crossing *eif4e1b* heterozygous embryos expressing 3xflag-sfGFP-eIF4E1B.

We also generated *ziwi:EGFP* transgenic fish by injecting a plasmid containing the *ziwi* promoter sequence driving *EGFP* (Leu & Draper, 2010). Juvenile fish expressing GFP in the gonads were selected as founders and crossed with *eif4e1b* homozygous fish. Wild-type and homozygous *eif4e1b* siblings containing the *ziwi:EGFP* transgene were further obtained by incrossing and genotyping.

### Histology of adult zebrafish ovaries

Ovaries from 7-month old wild-type and *eif4e1b* knockout females (3 fish per genotype) were dissected under the scope and fixed in 3.7% paraformaldehyde (PFA) in PBS overnight at 4 °C. Embryos were washed in PBS and embedded in 2% agarose followed by dehydration and paraffin infiltration in an automated tissue processor (Donatello Series 1, DiaPath). The processed agarose blocks were then embedded in paraffin using an embedding station (Tissue-Tek TEC, Sakura), and sectioned at 2 µm on a microtome Microm HM 355 S Leica (ThermoFisher). The sections were dried at 50 °C overnight. Prior to staining procedures, the slides were dewaxed in Thermo Scientific Shandon Xylene Substitute and further rehydrated through decreasing ethanol series using an automatic stainer (Epredia Gemini AS).

Hematoxylin and eosin (H&E) staining was also performed using Epredia Gemini AS. For phospho-histone H3 (phH3) staining, the slides were incubated for 30 min at 100 °C in EDTA retrieval solution. After cooling, slides were washed in TBS and incubated in the dark for 10 min with 3% H_2_O_2_. Slides were then washed with TBS and transferred to TBST. After creating a hydrophobic barrier with a PAP Pen (Abcam), the slides were incubated with 5 % BSA in TBST supplemented with 10% goat serum (Merck) for 1 h at room temperature (RT). Next, the slides were stained for 1 h with phH3 (06-570, Merck) at RT. After washing 3 times with TBST, slides were incubated with the polymer rabbit detection system (DCS) and phH3 staining was detected after incubating the slides with DAB substrate kit (Abcam) for 10 min at RT. The slides were counterstained with Epredia Shandon Harris Hematoxylin (Fisher Scientific) and further dehydrated using the Epredia Gemini AS. All slides were air dried overnight and covered with Eukitt Neo medium for coverslipper (O. Kindler) using an automatic coverslipper (Tissue-TEK GLC, Sakura). Images were taken with a Slide Scanner Pannoramic 250 (software version 3.0.2.127553; scanner hardware ID P250F20J2101).

### Protein expression in *E. coli*

The coding sequences of eIF4Ea (34-215; Uniprot: A8E579), eIF4E1c (49-230; Uniprot: B8A6A1), eIF4E2 (52-236; Uniprot: B2GPF6), and eIF4E3 (44-224; Uniprot: Q66HY7), were amplified from cDNA obtained from zebrafish embryos. The coding sequences of mouse (63-244; Uniprot: Q3UTA9) and human (60-242; Uniprot: A6NMX2) eIF4E1B were obtained as synthetic genes from Twist Bioscience. eIF4E coding sequences were cloned into a pOPINB vector providing an N-terminal 10xHis tag followed by an MBP tag and a 3C protease cleavage site. Cloning was done via Gibson assembly and specific mutations and deletions were introduced via site-directed mutagenesis. Plasmids containing the eIF4E-binding motifs of human eIF4G (608-647), eIF4EBP1 (50-83) and eIF4ENIF1 (27-63) with N-terminal MBP and C-terminal GB1 tags were obtained from C. Igreja (MPI for Biology, Tübingen) (eIF4G and eIF4EBP1 plasmids were previously published in Peter *et al*, 2015; Grüner *et al*, 2016). Plasmids were transformed in *E. coli* BL21 (DE3) cells. For protein expression, cells were grown in LB medium supplemented with antibiotic at 37 °C until reaching an OD_600_ in between 0.6 and 1. Cultures were induced with 0.25 mM IPTG (ThermoFisher) and grown overnight at 18 °C. Pellets were collected by centrifugation at 3,900 xg for 20 min.

### Pulldowns

Bacteria pellets were resuspended in cold lysis buffer containing 50 mM Tris-HCl pH 7.5, 100 mM NaCl, 1 mM DTT (Merck), 10 μg/mL DNase I (Merck) and cOmplete, EDTA-free protease inhibitor cocktail (Merck). Cells were lysed by sonication and lysates were collected via centrifugation at 21,000 xg for 20 min.

To test for protein-protein interactions, the supernatants (soluble fractions) were supplemented with imidazole (Merck) to reach a final concentration of 20 mM. HisPur Ni-NTA magnetic beads, previously equilibrated with washing buffer containing 50 mM Tris-HCl pH 7.5, 100 mM NaCl, 1 mM DTT (Merck), 20 mM imdidazole (Merck) and 0.01% Tween-20 (Merck), were incubated for 1 h with either the soluble fraction of the lysates containing His-tagged proteins (eIF4E or Lsm14^[5-79]^) or with lysis buffer at 4 °C. After 1 h, lysates were discarded, and beads were incubated with the soluble fraction of the putative interactors for 1 h at 4 °C. Beads were washed three times with washing buffer and bound proteins were eluted by incubating with the beads with elution buffer, containing 50 mM Tris-HCl pH 7.5, 100 mM NaCl, 1 mM DTT (Merck) and 500 mM imidazole (Merck), for 15 min at RT. The elution step was performed twice, and eluates were combined in one tube.

To test for mRNA cap binding, γ-aminophenyl-m^7^GTP (C10-spacer) agarose beads (Jena Bioscience) were packed into Pierce Screw Cap Spin Columns (ThermoFisher) and equilibrated with washing buffer containing 50 mM Tris-HCl pH 7.5, 100 mM NaCl, 1 mM DTT (Merck) and 0.01% Tween-20 (Merck). The soluble fractions of *E. coli* lysates were added to the columns and incubated for 5 min at RT before centrifugation at 2,000 xg for 30 sec. After repeating this step twice, beads were washed three times with washing buffer and incubated for 10 min with 2x Laemmli-sample buffer (Bio-Rad). The elution step was performed twice, and eluates were combined in one tube.

Boiled samples were analyzed by SDS-PAGE using Any kD precast Polyacrylamide Gels (Bio-Rad). Gels were stained with InstantBlue Coomassie Protein Stain (Abcam). Raw images of the gels were quantified with Fiji. Graphs and statistical analysis were done with GraphPad Prism 8.0.2.

### AlphaFold predictions and structural analysis

Structural predictions of protein complexes were performed with ColabFold (for the generation of multiple sequence alignments) and AlphaFold-Multimer (for the generation of structures) (Mirdita *et al*, 2022; Evans *et al*, 2022). Structures were visualized with ChimeraX-1.2.5 or Chimera 1.13.1. Models were aligned using the command *mmaker*.

### mRNA injections into zebrafish eggs

The coding sequences of full length *Dcp2* (Uniprot: Q6NYI8), *Ddx6* (Uniprot: E7FD91), *Ybx1* (Uniprot: B5DE31-2), *eIF4E1b* (Uniprot: Q9PW28), and *eIF4Ea* (see above) were PCR-amplified from zebrafish cDNA and cloned via Gibson assembly into a vector containing SP6 and T3 promoters, the 5’ and 3’ UTRs of Xenopus *β-globin* and an A^29^, C^14^ tail. P-body markers contained N-terminal *3xflag* and *dsRed* tags spaced by linkers, whereas *eIF4E1b* and *eIF4Ea* were tagged with N-terminal *3xflag* and *sfGFP* tags spaced by linkers. Specific mutations in *eIF4E1b* were introduced via site-directed mutagenesis. Capped mRNAs were transcribed from linearized plasmids using SP6 or T3 mMessanger Machine Kit (Ambion), according to the manufacturer’s protocol. mRNAs were purified with the RNA Clean & Concentrator kit (Zymo Research). For all injections, 100 pg of mRNAs were injected into one-cell stage embryos. Nuclei were labeled with *histone 2B (H2B)-RFP* mRNA (Stock *et al*, 2022).

### Microscopy

For confocal imaging, oocytes or manually dechorionated embryos were mounted in a drop of 0.8% low-melting agarose in PBS on round glass bottom dishes (Ibidi). For determining the localization of eIF4E proteins at specific stages of oogenesis and embryogenesis, samples were fixed in 3.7% paraformaldehyde (PFA) in PBS overnight at 4 °C. Samples were then washed in PBS and permeabilized with PBS-0.5% Triton X-100 for 30 min at RT; for nuclei or mitochondria staining, samples were incubated with DAPI (Sigma; 1:5000 dilution, 15 min incubation at RT) or Mitotracker Deep Red FM (Invitrogen; 1:2000 dilution, 30 min incubation at RT), respectively. Dishes were filled with E3 medium (5 mM NaCl, 0.17 mM KCl, 0.33 mM CaCl_2_, 0.33 mM MgSO_4_, 10^−5^% methylene blue). Images were acquired on an inverted LSM800 Axio Observer (Zeiss) with a temperature of incubation of 27°C for live imaging. The following objectives were used: Plan-Neofluor 10x/0.3 (Zeiss), Plan-Apochromat 20x/0.8 M27 (Zeiss) or Plan-Apochromat 63x/1.2 water objective (Zeiss). Images from different fluorescent channels were acquired separately with a GaAsP-Pmt detector using different excitation (ex) and detection (em) wavelengths depending on the fluorophore used (ex/em): dsRed (561/563-617 nm), sfGFP (488/400-527, 502-700 or 410-521 nm depending on the experiment), mRFP1 (561/564-617 nm), farRed (640/656-700 nm) or DAPI (405/410-456 nm). ZenBlue 3.1 (Zeiss) was used for image acquisition. For granule quantification of mRNA injections, 2D-images were taken with the Plan-Apochromat 63x/1.2 water objective (Zeiss). For detecting and quantifying the granules, we manually annotated a training dataset and created a custom tensorflow model using CSBDeep/CARE, which we then applied to our images via Fiji. A threshold was set on the resulting images for subsequent segmentation and the resulting regions were counted and measured.

For fluorescent widefield microscopy, juvenile fish of 1-2 cm in length were anesthetized in 0.1% (w/v) tricaine (Sigma). Images were acquired on a fluorescent stereomicorcope (Lumar.V12, Zeiss) with a NeoLumar S 0.8x objective (Zeiss), an Axiocam 702mono (Zeiss) and an X-cite Xylis (Exelitas) fluorescent lamp using ZenBlue 3.1 (Zeiss) software.

Images were post-processed with ImageJ 1.53t software adjusting brightness and contrast.

### Immunoprecipitation followed by mass spectrometry (IP-MS)

Embryos were dechorionated with 1 mg/mL of pronase (Sigma-Aldrich), lysed in cold lysis buffer containing 100 mM Tris-HCL pH 7.5, 150mM NaCl, 0.05% Triton X-100 (Sigma-Aldrich) and cOmplete, EDTA-free protease inhibitor cocktail (Merck) and homogenized with a Dounce homogenizer (Sigma-Aldrich). Lysates were cleared at 13000 xg for 10 min at 4 °C and incubated with anti-GFP agarose beads (Molecular Biology Service, IMP), which were previously equilibrated with lysis buffer overnight at 4 °C. Beads were then washed ten times with washing buffer containing 100 mM Tris-HCl pH 7.5 and 150 mM NaCl. For RNAse treatment, lysates were incubated with 1 unit of RNAse I (Ambion) per 500 μL lysate for 30 min at 37 °C. For immunoprecipitation on squeezed eggs, mature oocytes were squeezed from two females into Petri dishes and immediately lysed and homogenized. For immunoprecipitation on oocytes, ovaries from three female fish were pooled. Early (stages I and II) and late (stages III and IV) oocytes were harvested according to Jamieson-Lucy & Mullins, 2019 with some modifications. Briefly, the pooled ovaries were kept in Leibovitz’s L-15 medium (Gibco) at 28 °C. Ovaries were incubated in digestive mix containing 1 mg/ml collagenase I (Sigma Aldrich), 1 mg/ml collagenase II (Sigma Aldrich) and 0.5 mg/ml hyaluronidase IV (Sigma-Aldrich) in Leibovitz’s L-15 medium for 30 min at RT on a tube rotator. After stopping the digest with Leibovitz’s L-15 medium, oocytes were manually dissociated, washed with Leibovitz’s L-15 medium and lysed and homogenized as described before.

HPLC-MS was performed using an UltiMate 3000 RSLC nano system coupled to an Orbitrap Exploris 480 mass spectrometer, equipped with a Proxeon nanospray source or to an Orbitrap Eclipse Tribrid mass spectrometer equipped with a FAIMS pro interface and a Nanospray Flex ion source (all parts Thermo Fisher Scientific). Peptides were loaded onto a trap column (Thermo Fisher Scientific, PepMap C18, 5 mm × 300 μm ID, 5 μm particles, 100 Å pore size) at a flow rate of 25 μL/min using 0.1% TFA as mobile phase. After 10 min, the trap column was switched in line with the analytical column (Thermo Fisher Scientific, PepMap C18, 500 mm × 75 μm ID, 2 μm, 100 Å). Peptides were eluted using a flow rate of 230 nl/min, starting with the mobile phases 98% A (0.1% formic acid in water) and 2% B (80% acetonitrile, 0.1% formic acid) and linearly increasing to 35% B over the next 120 min, followed by a gradient to 95% B in 5 min, staying there for 5 min and decreasing in 2 min back to the gradient 98% A and 2% B for equilibration at 30°C. The Orbitrap Exploris 480 mass spectrometer was operated in data-dependent mode, performing a full scan (m/z range 350-1200, resolution 60,000, normalized AGC target 100%) at 3 different compensation voltages (CV -45, -60, -75), followed each by MS/MS scans of the most abundant ions for a cycle time of 0.9 (CV -45, -60) or 0.7 (CV -75) seconds per CV. MS/MS spectra were acquired using HCD collision energy of 30, isolation width of 1.0 m/z, orbitrap resolution of 30,000, normalized AGC target 200%, minimum intensity of 25,000 and maximum injection time of 100 ms. Precursor ions selected for fragmentation (include charge state 2-6) were excluded for 45 s. The monoisotopic precursor selection (MIPS) filter and exclude isotopes feature were enabled. The Eclipse was operated in data-dependent mode, performing a full scan (m/z range 375-1500, resolution 60k, AGC target value 400,000, normalized AGC target 100%) at 3 different compensation voltages (CV-45, -60, -75), followed each by MS/MS scans of the most abundant ions for a cycle time of 0.9 sec per CV. MS/MS spectra were acquired using an isolation width of 1.0 m/z, Orbitrap resolution of 30,000, AGC target value of 100,000, intensity threshold of 25,000 and a maximum injection time of 50 ms, using the Orbitrap for detection with HCD collision energy of 30. Precursor ions selected for fragmentation (include charge state 2-6) were excluded for 45 s. The monoisotopic precursor selection filter and exclude isotopes feature were enabled. Raw MS data from IP experiments was loaded into Proteome Discoverer (PD, version 2.5.0.400, Thermo Scientific). All MS/MS spectra were searched using MS Amanda v2.0.0.19924 (Dorfer *et al*, 2014). Trypsin was specified as a proteolytic enzyme cleaving after lysine and arginine (K and R) without proline restriction, allowing for up to 2 missed cleavages. Mass tolerances were set to ±10 ppm at the precursor and ±10 ppm at the fragment mass level, increasing to ±20 ppm for the Eclipse data. Peptide and protein identification was performed in two steps. An initial search was performed against the ENSEMBL database, using taxonomy zebrafish (release 2021_06; 45,694 sequences; 26,805,620 residues), with common contaminants appended. Here, beta-methylthiolation of cysteine was searched as fixed modification, whereas oxidation of methionine, deamidation of asparagine and glutamine and glutamine to pyro-glutamate conversion at peptide N-termini were defined as variable modifications. Results were filtered for a minimum peptide length of 7 amino acids and 1% FDR at the peptide spectrum match (PSM) and the protein level using the Percolator algorithm (Käll *et al*, 2007) integrated in Proteome Discoverer. Additionally, an Amanda score of at least 150 was required. Identified proteins were exported and subjected to a second step search considering phosphorylation of serines, threonines and tyrosines as additional variable modifications. The localization of the post-translational modification sites within the peptides was performed with the tool ptmRS, based on the tool phosphoRS (Taus *et al*, 2011). Identifications were filtered using the filtering criteria described above, including an additional minimum PSM-count per protein in at least one sample of 2. The identifications were subjected to label-free quantification using IMP-apQuant (Doblmann *et al*, 2019). Proteins were quantified by summing unique and razor peptides and applying intensity-based absolute quantification (iBAQ) (Schwanhäusser *et al*, 2011). Protein-abundances-normalization was done using sum normalization. Statistical significance of differentially expressed proteins was determined using limma (Smyth, 2005).

### Western blotting

Lysates from early oocytes (stages I-II) were supplemented with 4x laemmli-sample buffer (Bio-Rad). 4 hpf embryos were deyolked in 55 mM NaCl, 1.8 mM KCl, 1.25 mM NaHCO_3_, and cOmplete, EDTA-free protease inhibitor cocktail (Merck). Embryos were lysed by pipetting and the yolk was dissociated by shaking the lysates at 1100 rpm for 5 min at 4 °C. Cells were pelleted by centrifugation at 500 rpm for 30 s and resuspended in 4x laemmli-sample buffer (Bio-Rad). Oocyte and embryo samples were boiled at 95 °C for 5 min and separated by SDS-PAGE using Any kD precast Polyacrylamide Gels (Bio-Rad). Blotting was performed onto a nitrocellulose membrane (GE Healthcare) using a Bio-Rad wet blot system. Membranes were blocked for 1 h at RT with 3% BSA in TBST, and the following antibodies were added to a solution containing 1.5% BSA in TBST: anti-Dcp2 (rabbit, 1:1000, NBP2-16109, Novus Biological), which was previously tested in zebrafish embryos (Zampedri *et al*, 2016) and anti-Rps17 (rabbit, 1:1000, ab128671, Abcam). After washing with TBST, membranes were incubated with F(ab′)2 anti-rabbit IgG (H+L)-HRPO secondary antibody (goat, 1:10000, 111-036-045, Dianova). Chemiluminescence was induced by using Clarity Western ECL Substrate (Bio-Rad).

### RNA immunoprecipitation (RIP)

8-cell stage embryos were dechorionated with 1 mg/mL of pronase. RIP was performed as described in Ren *et al*, 2020 with some modifications. Briefly, embryos were lysed in cold RIP buffer containing 50 mM Tris-HCl pH 7.5, 150 mM NaCl, 5 mM MgCl_2_, 0.5% IGEPAL^®^ CA-630 (Sigma-Aldrich), 1 mM DTT (Merck), cOmplete, EDTA-free protease inhibitor cocktail (Merck) and 40 U/μL RNAse Out (Invitrogen) and homogenized with a Dounce homogenizer (Sigma-Aldrich). Lysates were cleared at 13000 xg for 10 min at 4 °C and incubated with anti-GFP agarose beads (Molecular Biology Service, IMP), previously equilibrated with RIP-buffer for 2 h at 4 °C. 200 μL of each lysate, corresponding to the input sample, was kept on ice. Beads were washed two times with RIP buffer and two times with washing buffer containing 50 mM Tris-HCl pH 7.5, 150 mM NaCl, 5 mM MgCl_2_ and 0.5% IGEPAL^®^ CA-630 (Sigma-Aldrich). Beads were incubated with 0.3 μg/μl proteinase K, 50 mM Tris-HCL pH 7.5, 150 mM NaCl, 5 mM MgCl_2_, 0.5% IGEPAL^®^ CA-630 (Sigma-Aldrich) and 1% SDS for 30 min at 55 °C. RNA from elution and input samples was purified with the RNA Clean & Concentrator kit (Zymo Research) according to the manufacturer’s protocol.

### RNA-seq

Samples were submitted to Macrogen for library preparation and NGS sequencing. Input samples (corresponding to total RNA isolated from embryo lysates) were prepared with a SMARTer Stranded RNA library (Ribo-Zero) protocol, whereas RIP samples were prepared with SMARTer Ultra Low RNA kit. RNA samples were sequenced using NovaSeq 6000 platform (Illumina). Reads were aligned to the zebrafish genome (GRCz11) and differential expression analysis was performed with the DESeq2 package (Love *et al*, 2014) in R 3.4.1. Graphs of DEG analyses were generated in RStudio 2021.09.2; genes encoding proteins belonging to the following InterPro families were highlighted in **Fig 5A**: IPR009072 (histone-fold), IPR005819 (linker histone H1/H5), IPR001951 (histone H4), IPR019809 (histone H4, conserved site), IPR035425 (CENP-T/histone H4, histone fold), IPR000164 (histone H3/CENP-A), IPR007125 (histone H2A/H2B/H3), IPR000558 (histone H2B), IPR002119 (histone H2A), IPR032454 (histone H2A, C-terminal domain), IPR032458 (histone H2A conserved site), IPR005818 (linker histone H1/H5, domain H15), IPR021171 (core histone macro-H2A), and IPR035796 (core histone macro-H2A, macro domain). Reads were visualized with Integrative Genome Viewer (IGV) version 2.11.9. Comparisons between the data generated in this study with previously published datasets were done in Microsoft Excel version 16.51. Graphs were generated in GraphPad Prism 8.0.2.

### Data availability

Previously published structures used in this study are available at RCSB Protein Data Bank (RCSB PDB): 5BXV, 5T46, 4UE9. Translational efficiency and polyA tail length data were previously published and accessible at Gene Expression Omnibus (GEO; accession number: GSE52809) and at https://zenodo.org (names: hs25.h5.xz.part01 to 11), respectively. Available RNA-seq data from different stages of zebrafish embryogenesis and adult tissues: GSE147112 (oogenesis and embryogenesis), GSE3290031 (subseries GSE32898; embryogenesis), GSE111882 (ovaries), and GSE171906 (adult tissues and organs). RIP-seq and RNA-seq data generated in this study has been submitted to GEO, with the accession number GSE233570. Mass spectrometry proteomics data has been deposited to the ProteomeXchange Consortium via the PRIDE (Perez-Riverol *et al*, 2019) partner repository with the dataset identifier PXD042434.

## Acknowledgements

We thank A. Schleiffer (Bioinformatics, IMP) for performing the phylogenetic analysis of eIF4E genes across vertebrates; M. Novatchkova (Bioinformatics, IMP) and J. Elbers for their help with RIP-seq data analysis; P. Bhat for analyzing the TAIL-seq data; C. Igreja (MPI for Biology, Tübingen) for providing the plasmids for the expression of the human eIF4E-binding motifs of eIF4G, eIF4EBP1 and 4E-T in *E. coli,*; B. Draper (UC Davis) for providing the *ziwi:EGFP* plasmid for transgenesis; the Proteomics facility at the Vienna BioCenter Core Facilities (VBCF), in particular O. Hudecz, R. Imre, S. Opravil, K. Mechtler and E. Roitinger, for the processing, analyses and supervision of MS data; the Histology facility at VBCF, in particular L. Ushakova and M. Grivej, for the staining of zebrafish ovaries; the BioOptics facility at IMP, in particular T. Lendl, P. Pasierbek and A. Moreno-Cencerrado, for their help with microscopy data acquisition and analysis; the IMP animal facility, especially F. Puhl, K. Rattner, J. König and D. Sunjic for their care of zebrafish; T. Mylenko, A. Bandura, G. Deneke, and M. Binner for their help with genotyping; the Molecular Biology Service at IMP for Sanger sequencing and for providing competent cells and reagents; J. Ahel for implementing AF Multimer in the VBC cluster; the entire Pauli lab and the VBC RNA Salon for critical feedback and valuable discussions; U. Hohmann, J. Roehsner, A. Chugunova, L. Veleti and C. Plaschka for feedback on the manuscript. Research in the lab of AP was funded by the Research Institute of Molecular Pathology (IMP), Boehringer Ingelheim, the Austrian Academy of Sciences, FFG (Headquarter grant FFG-852936), the FWF START program (Y 1031-B28), the ERC Consolidator grant 101044495/GaMe, an HFSP Career Development Award (CDA00066/2015), an HFSP Young Investigator Grant (RGY0079/2020) and the FWF SFB RNADeco (project number F80). L.L.-O. was supported by an SNF Early Postdoc.Mobility fellowship (P2GEP3_191204), an EMBO long-term fellowship (ALTF 1165-2019) and an MSCA-IF-EF-SE (890218).

## Author contributions

**Laura Lorenzo-Orts**: conceptualization, data curation, formal analysis, funding acquisition, investigation, methodology, project administration, supervision, validation, visualization, writing – original draft. **Marcus Strobl**: data curation, investigation, methodology, validation, visualization. **Benjamin Steinmetz**: data curation, formal analysis, investigation, methodology. **Friederike Leesch**: methodology. **Carina Pribitzer**: methodology. **Michael Schutzbier**: methodology, resources. **Gerhard Dürnberger**: data curation, formal analysis, resources, supervision. **Andrea Pauli**: conceptualization, data curation, funding acquisition, project administration, resources, supervision, writing – original draft.

## Disclosure and competing interests statement

The authors declare that they have no conflict of interest.

---

**Figure EV1.**
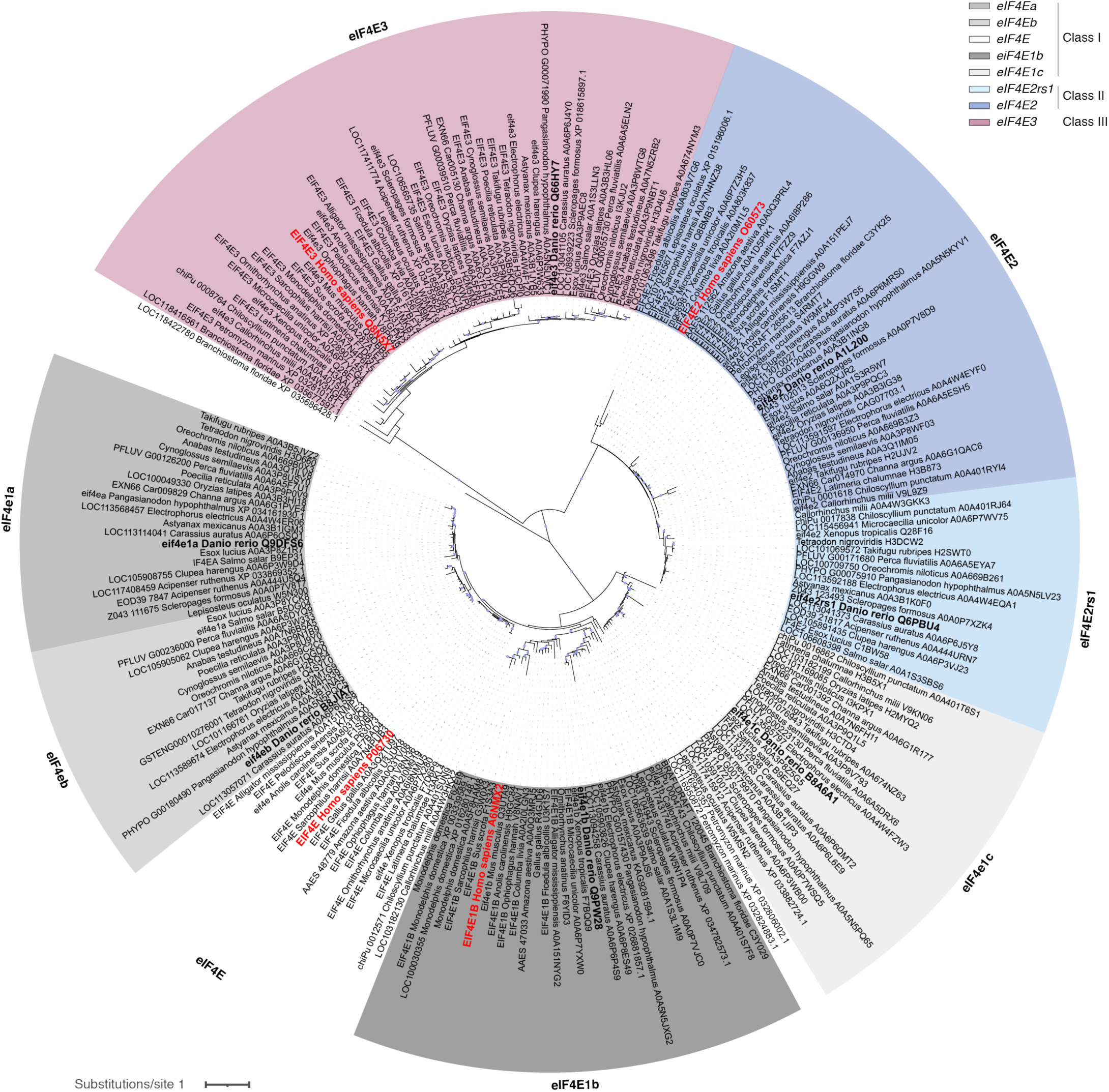
Phylogenetic analysis of vertebrate eIF4Es. A phylogenetic tree shows 3 major branches that correspond to 3 eIF4E classes: class I with eIF41a/eIF4Ea, eIF4Eb, eIF4E, eIF4E1b and eIF4E1c, class II with eIF4E2rs1and eIF4E2, and class III with eIF4E3. Branches that are supported by an ultrafast bootstrap (UFBoot) value ≥ 95% are indicated by a grey dot. Branch lengths represent the inferred number of amino acid substitutions per site, and branch labels are composed of gene name (if available), genus, species, and accession number. Zebrafish genes are bold black, human genes are bold red.

**Figure EV2.**
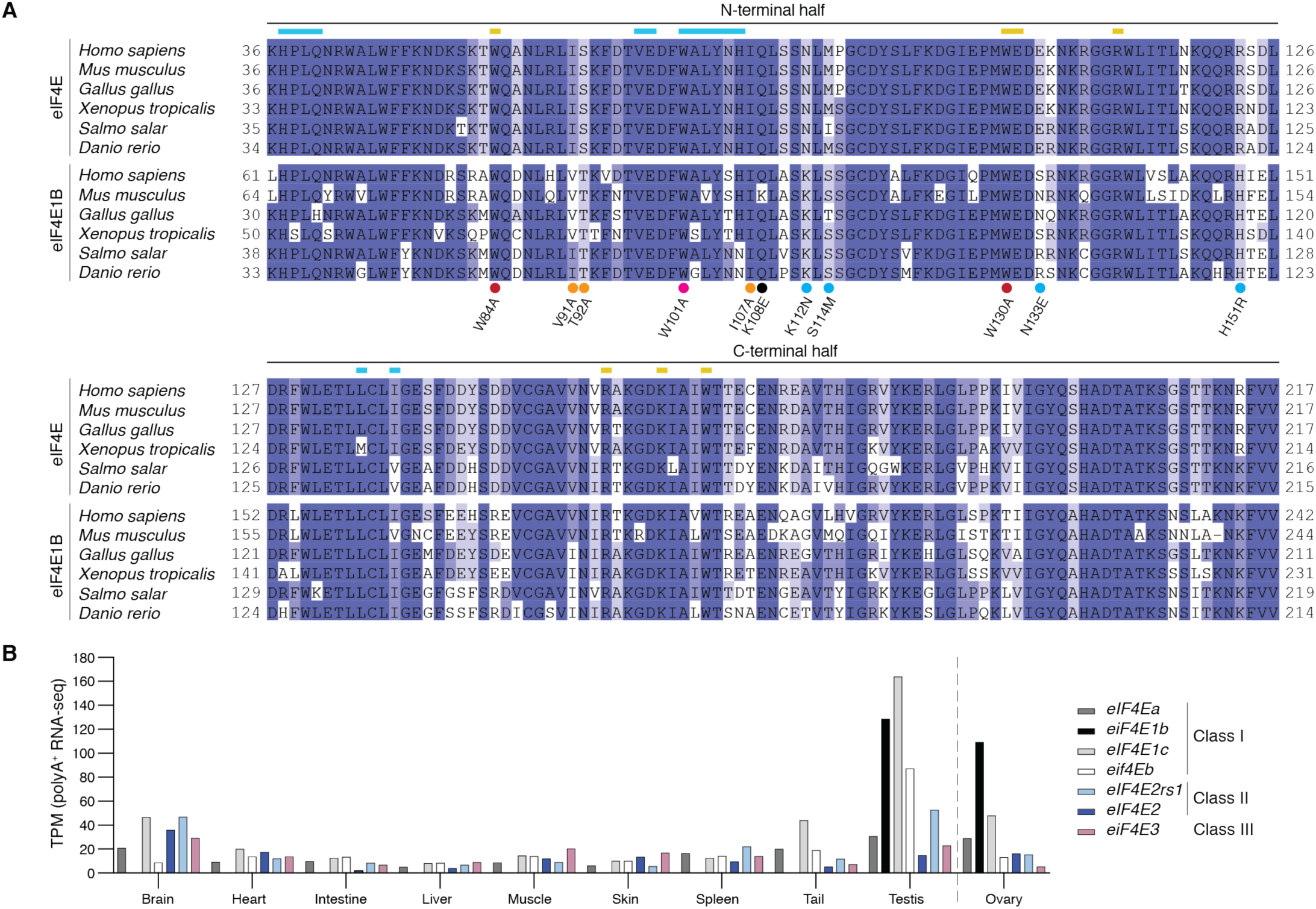
eIF4E1b and eIF4E are highly conserved but have specific expression patterns. **A** Amino acid alignment of eIF4E and eIF4E1B proteins from six vertebrate species. Unstructured N-terminal regions (see **Fig 1A**), which are not included in the constructs used for expressing recombinant proteins, are excluded from the alignment. Residues interacting with the mRNA cap or with eIF4E-binding motifs are highlighted with yellow or blue lines, respectively. eIF4E1B residues mutated in **Fig 3D** are indicated with dots (red: mRNA cap binding; pink: dorsal site; orange: lateral site; blue: conserved in eIF4Es but different in eIF4E1Bs; black: others). N-terminal and C-terminal regions exchanged in chimeric constructs are indicated. **B** mRNA levels (in transcripts per million, TPM) of zebrafish eIF4Es in different organs and adult tissues based on polyadenine-selected RNA-seq data (Fujihara *et al*, 2021) (ovary RNA-seq data from Herberg *et al*, 2018).

**Figure EV3.**
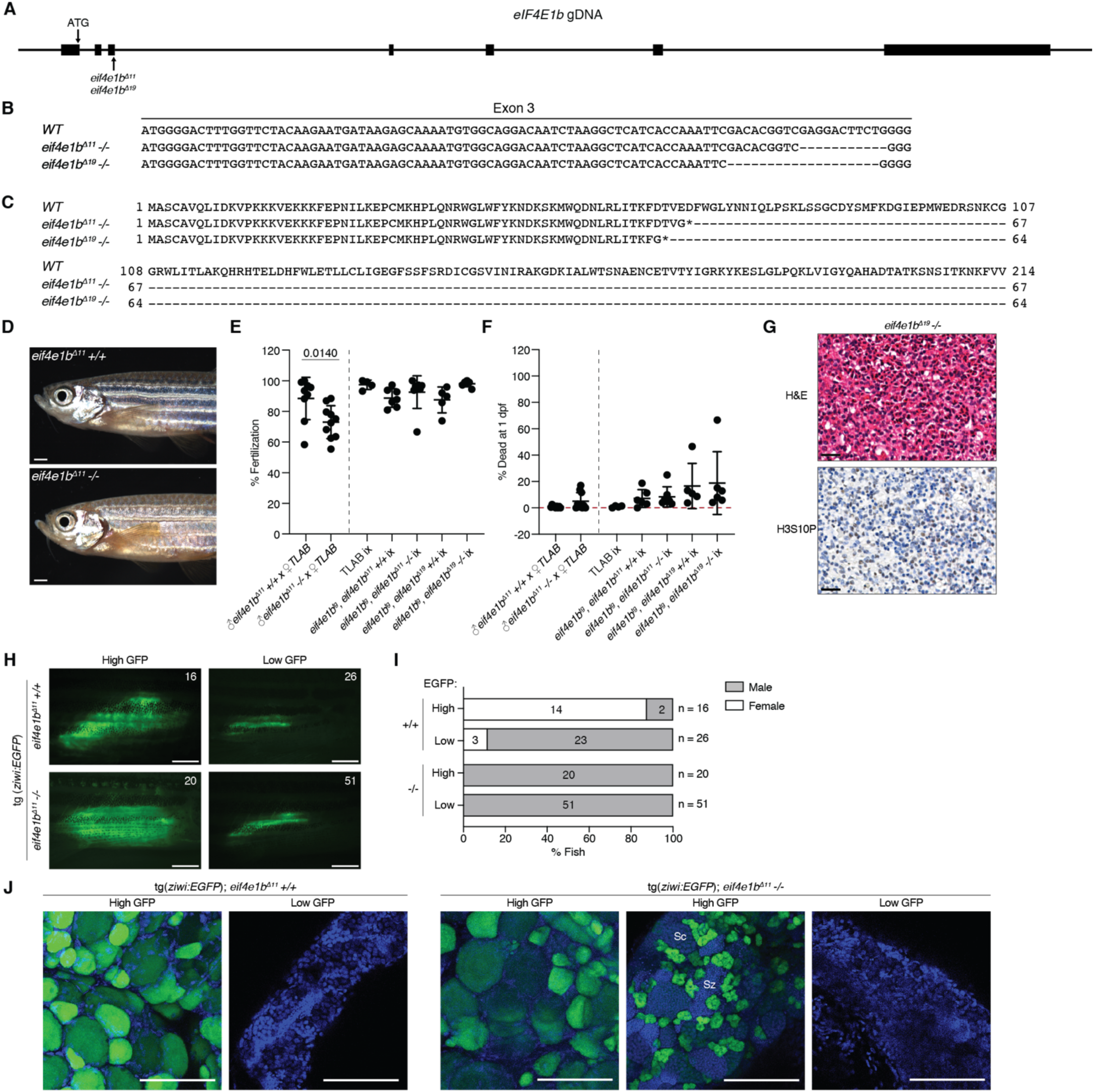
*eif4e1b* mutants develop into fertile males due to sex reversal. **A** Scheme of the zebrafish *eIF4E1b* locus, indicating the start codon (*ATG*) and the 11 and 19 nt deletions present in the two *eif4e1b* mutants (*eif4e1b^Δ11^*and *eif4e1b^Δ19^*, respectively) generated in this study. Exons are depicted as boxes, introns as lines. **B** Nucleotides missing in the third exon of *eif4e1b* mutants. **C** *eIF4E1b* mRNA translation in *eif4e1b* mutants result in truncated proteins of 67 (for *eif4e1b^Δ11^*) and 64 (for *eif4e1b^Δ19^*) amino acids. Asterisks indicate premature stop codons. **D** Representative pictures of *eif4e1b^Δ11^* mutant and wild-type sibling males obtained from a heterozygous incross (scale bars = 1 mm). **E** Fertilization rates of embryos obtained by crossing *eif4e1b^Δ11^* homozygous or wild-type males with wild-type females (left). Overexpression of *3xflag-sfGFP-eIF4E1b* (*eif4e1b^tg^*) in transgenic *eif4e1b* wild-type and mutant siblings results in fertile males and females (right; ix = incross). **F** Percentage of embryos dead at 1-day post fertilization (dpf). Embryos were obtained by crossing homozygous or wild-type *eif4e1b^Δ11^* male siblings with wild-type females (left), or by incrossing (ix) wild-type or mutant fish expressing 3xflag-sfGFP-eIF4E1b (right). **G** Hematoxylin and eosin (H&E, top) and phospho-histone H3 (bottom) staining of *eif4e1b ^Δ19^* ovaries. Scale bars = 20 μm. **H** Live microscopy of gonads from juvenile *eif4e1b^Δ11^* wild-type and mutant siblings expressing eGFP under the control of the *ziwi* promoter (scale bars = 1 mm). Fish were classified based on the area and intensity of eGFP. Numbers (top right) indicate the number of fish in each category. **I** Number of fish in *H* that developed into males or females. **J** Confocal microscopy of fixed gonads isolated from wild-type (left) and homozygous (right) *eif4e1b^Δ11^* juveniles expressing eGFP under the control of the *ziwi* promoter. Nuclei were stained with DAPI (in blue; Sc = spermatocytes; Sz = spermatozoa; scale bars = 100 μm). Data information: In *E* and *F*, significance for the first two genotypes was calculated using unpaired t-tests. The other genotypes were compared using one-way ANOVA followed by Tukey’s test (if not indicated, p-value > 0.005).

**Figure EV4.**
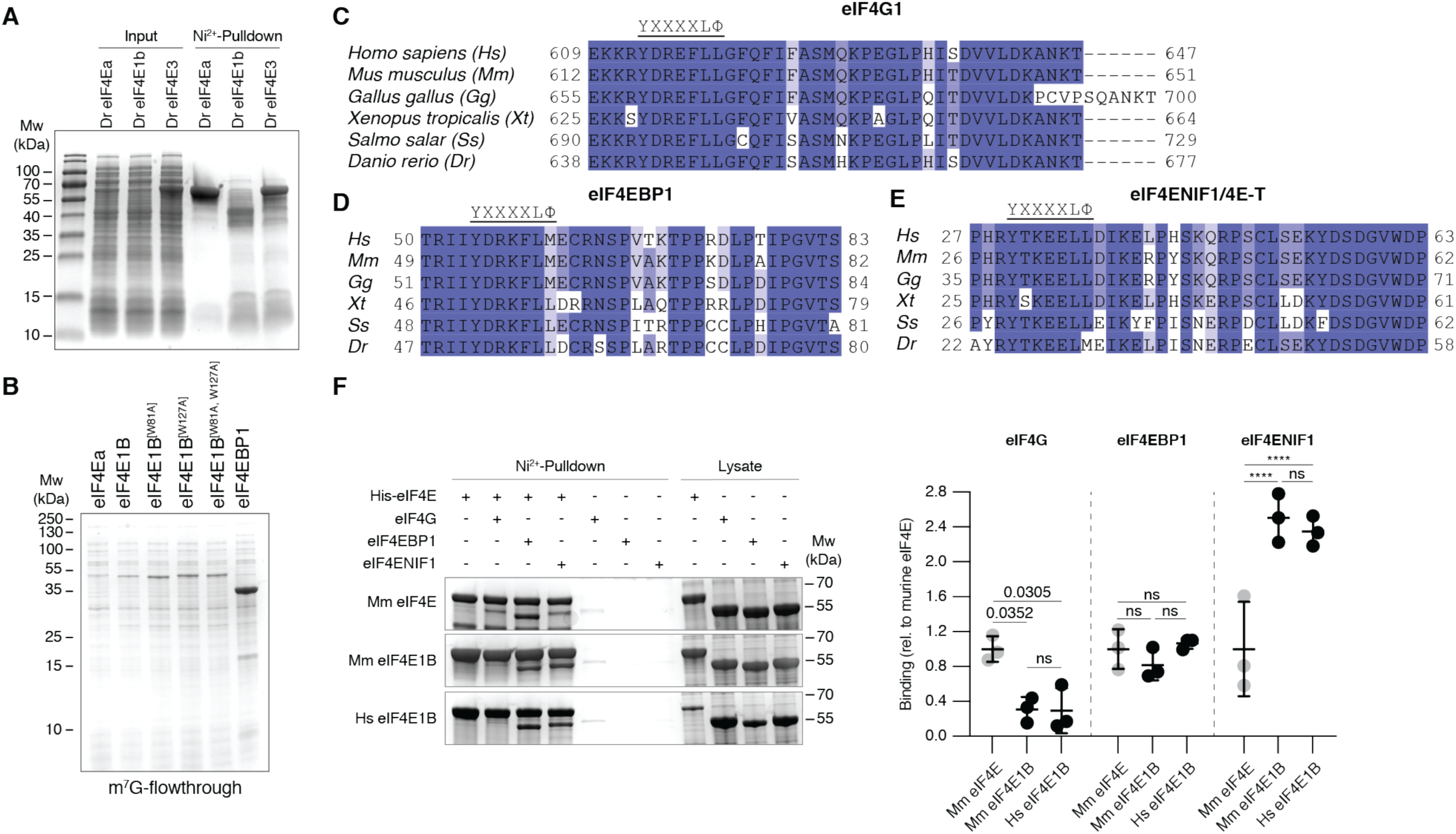
Human eIF4E1B interacts with the eIF4E-binding motifs of human eIF4EBP1 and eIF4ENIF1, which are conserved across vertebrates. **A** Coomassie stained gel of a pulldown assay using Ni^2+^ beads and bacterial lysates containing His and MBP tagged zebrafish (*Danio rerio*, Dr) eIF4Ea, eIF4E1b and eIF4E3. Inputs corresponding to the soluble fractions of the lysates are shown on the left, elutions are shown on the right. Predicted molecular weights (Mw) are 64.6 kDa for eIF4Ea and eIF4E1b and 64.1 kDa for eIF4E3. **B** Coomassie stained gel of flow-through fractions from an immunoprecipitation assay using m^7^G-coated beads and *E. coli* lysates containing eIF4Es and eIF4EBP1. **C-E** Amino acid alignments of the eIF4E-binding motifs of eIF4G1 (*B*), eIF4EBP1 (*C*) and eIF4ENIF1/4E-T (*D*) from six vertebrate species. The species abbreviations are shown in *C*. The canonical YXXXXLΦ eIF4E-binding motif is indicated (X = any amino acid; Φ = hydrophobic residue). **F** Human and mouse eIF4E1Bs show similar affinities for eIF4E-binding proteins. Representative Coomassie-stained gels of in vitro pulldown assays with bacteria lysates is shown on the left, quantification is shown on the right (n = 3 independent experiments; mouse eIF4E and eIF4E1B data is also shown in **Fig 2D**). Statistical analysis was performed with two-way ANOVA followed by Sidak’s multiple comparisons test (ns: non-significant).

**Figure EV5.**
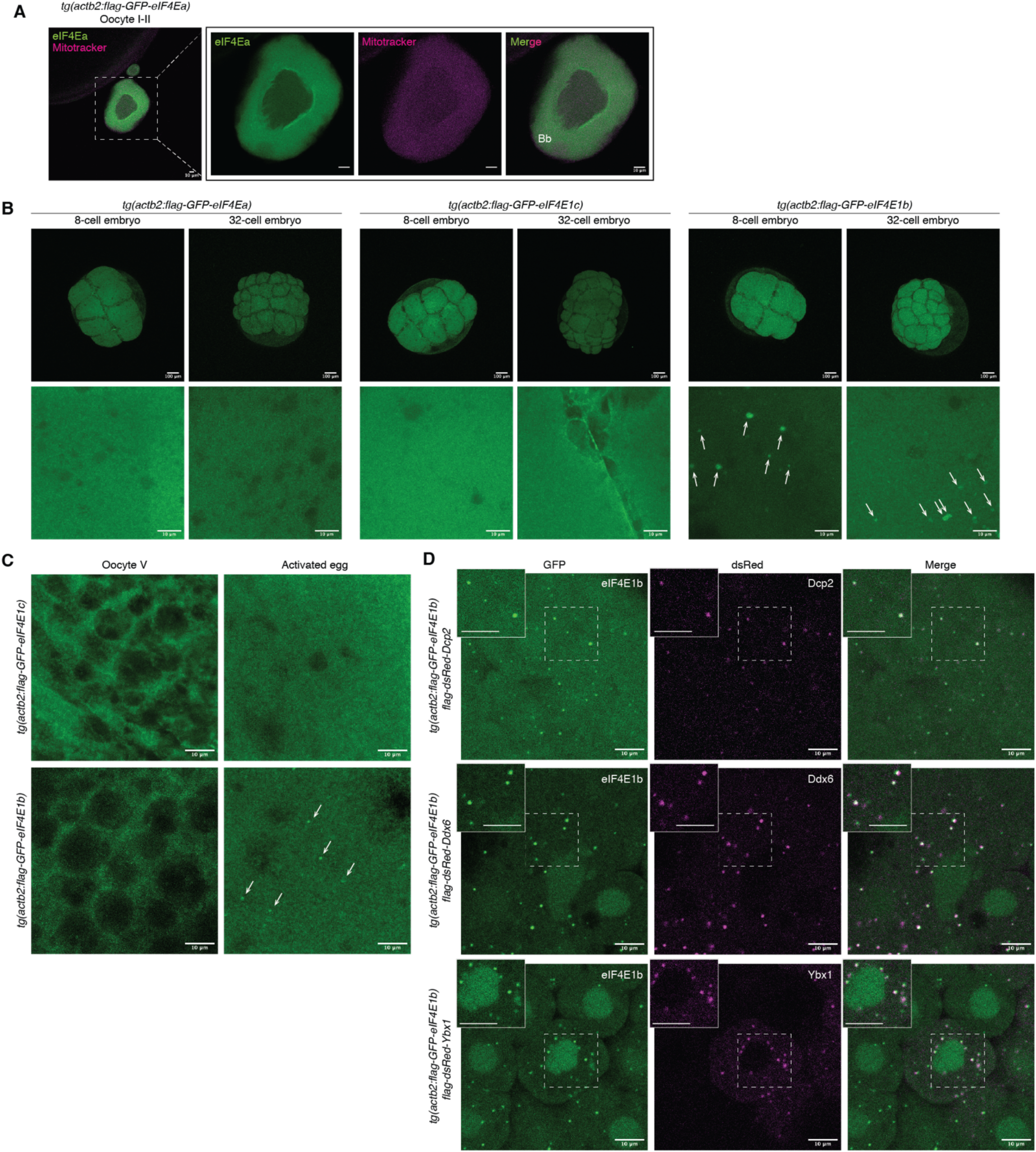
eIF4E1b localizes to P-bodies in the embryo. **A** eIF4Ea localizes to the cytosol of fixed early zebrafish oocytes. Mitochondria present in the Balbiani body (Bb) are stained with Mitotracker. Scale bars = 10 μm. **B** Confocal microscopy images of fixed transgenic embryos expressing 3xflag-sfGFP-tagged eIF4Ea, eIF4E1b and eIF4E1c (scale bars: top = 100 μm; bottom = 10 μm). eIF4E1b foci are indicated by white arrows. **C** Confocal microscopy images of squeezed (oocyte V) and activated eggs from transgenic lines expressing 3xflag-sfGFP-tagged eIF4E1c (top) and eIF4E1b (bottom). eIF4E1b foci are indicated by white arrows. **D** Colocalization experiments in zebrafish embryos expressing 3xflag-sfGFP-eIF4E1b (constitutively) and P-body markers (transiently). mRNAs containing the coding sequences of dsRed-tagged P-body components (Dcp2, Ddx6 and Ybx1) were injected into 1-cell embryos. Images were taken at 3 hours post fertilization. Regions enclosed by dashed boxes are shown at higher magnification on the top left (scale bars = 10 μm).

**Figure EV6.**
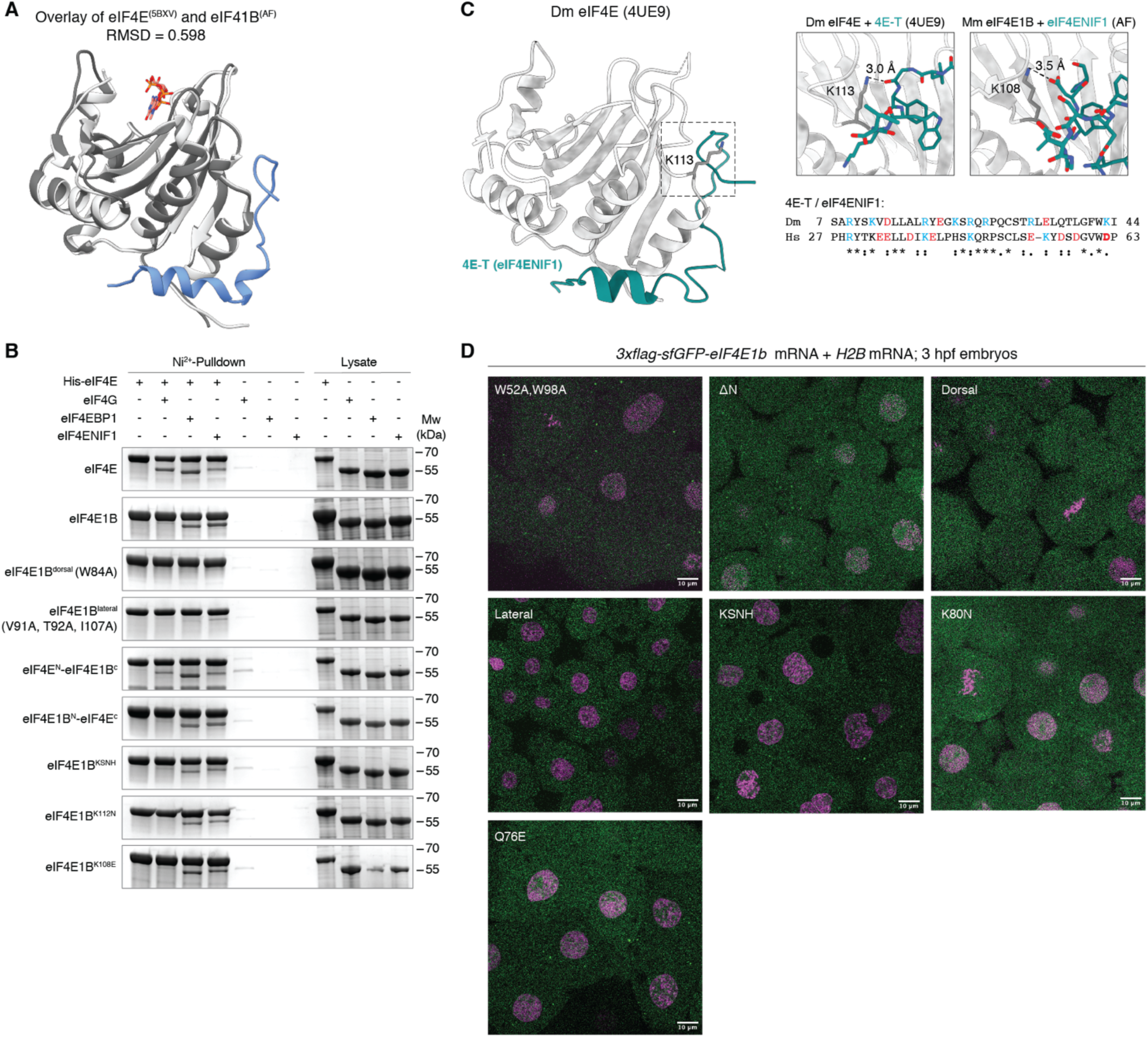
Coomassie-stained gels and confocal microscopy images of eIF4E1b mutants. **A** Superimposed structures of mouse eIF4E (PDB-5BXV, Sekiyama *et al*, 2015) and mouse eIF4E1B (predicted by AlphaFold, AF). RMSD = root-mean-square deviation. **B** Representative Coomassie-stained gels from pulldown assays with mouse eIF4E1B mutants. Quantifications are shown in **Fig 3D**. **C** Structural representation of *Drosophila melanogaster* (Dm) eIF4E in complex with 4E-T (PDB-4UE9, Peter *et al*, 2015). The nitrogen of Lys113 of Dm eIF4E (corresponding to Lys108 of mouse eIF4E1B) forms a main-chain contact with the carbonyl oxygen of Gly40 of Dm 4E-T. Although the eIF4E-binding motifs of Dm 4E-T and Hs eIF4ENIF1 are poorly conserved (Clustal Omega alignment on the bottom-right; * = identical residues; : = similar residues), Lys108 of mouse eIF4E1B is predicted to interact with Asp62 of eIF4ENIF1 in AF. Interactions are indicated by dashed lines. **D** Confocal microscopy images of 1k-cell embryos transiently expressing eIF4E1b mutant proteins from mRNAs injected at the 1-cell stage. Nuclei were labeled with *H2B-RFP* mRNA. Quantification of the number of GFP-positive foci is shown in **Fig 3H**. Scale bars = 10 μm.

**Figure EV7.**
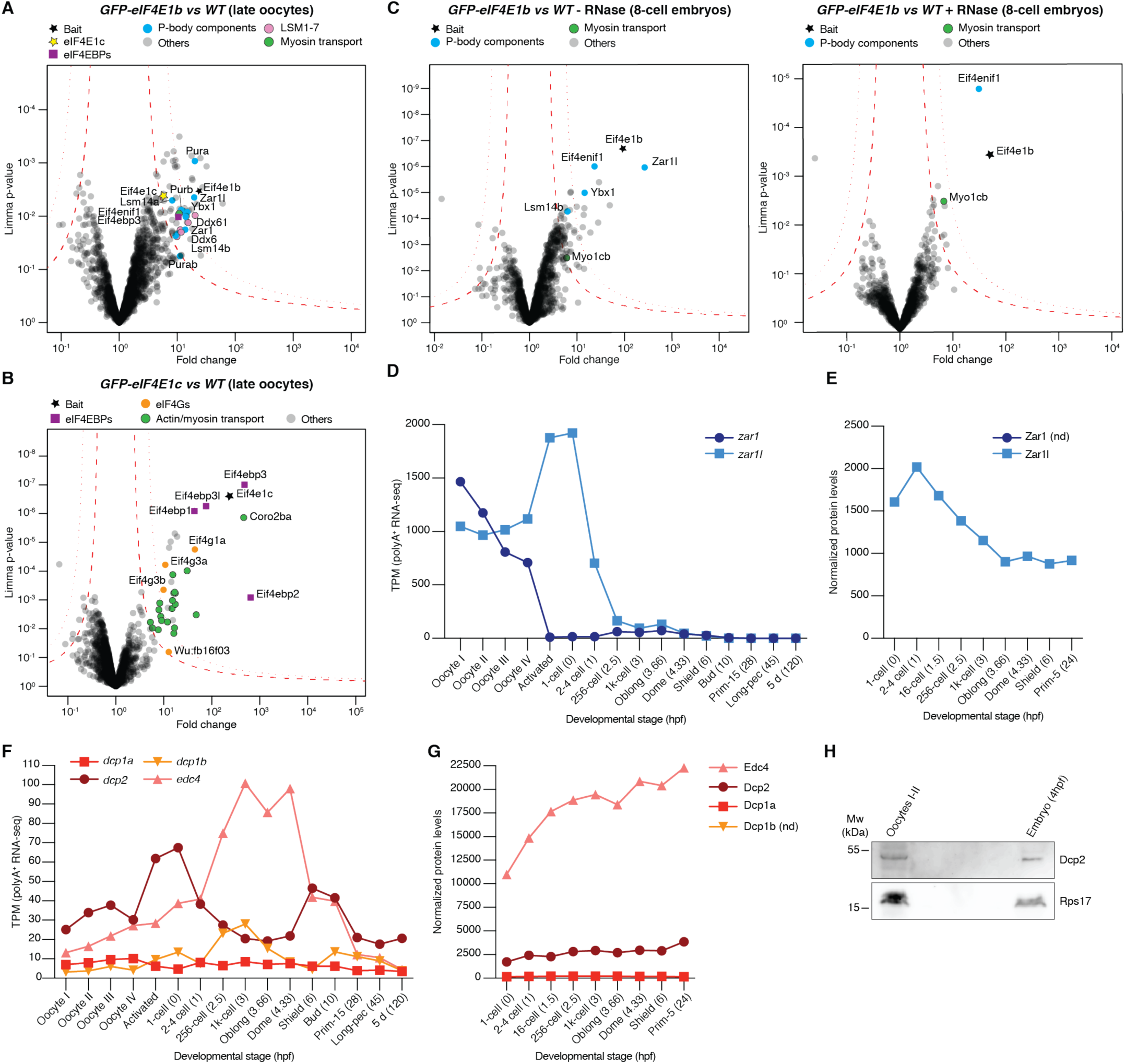
Additional IP-MS and expression data. **A-B** Volcano plots of the proteins identified by IP-MS in zebrafish late oocytes (stage III-IV) expressing GFP-tagged eIF4E1b (*A*) or eIF4E1c (*B*) compared to wild type (*WT*). **C** Volcano plots of the proteins identified by IP-MS in 8-cell embryos expressing GFP-tagged eIF4E1b compared to *WT* in the absence (mock, left) or presence (right) of RNase I. **D** Levels of *zar1* and *zar1l* mRNAs during zebrafish oogenesis and embryogenesis based on polyA-selected RNA-seq data (Pauli *et al*, 2012; Cabrera-Quio *et al*, 2021). **E** Normalized expression of Zar1l protein during zebrafish embryogenesis in TMT-MS. Zar1 peptides were not detected (nd). **F-G** mRNA (*F*) and protein (*G*) levels of decapping factors during zebrafish oogenesis and embryogenesis based on polyA-selected RNA-seq (Pauli *et al*, 2012; Cabrera-Quio *et al*, 2021) and TMT-MS data, respectively. **H** Western blots showing the expression of Dcp2 (predicted Mw of 45.5 kDa) and Rps17 (loading control, predicted Mw of 15.4 kDa) in zebrafish early oocytes and embryos. Data information: In *A-C*, permutation-based false discovery rates (FDRs) are displayed as dotted (FDR < 0.01) or dashed (FDR < 0.05) lines (n= 3 biological replicates). Hpf: hours post fertilization; Mw: molecular weight; TPM: transcripts per million.

**Figure EV8.**
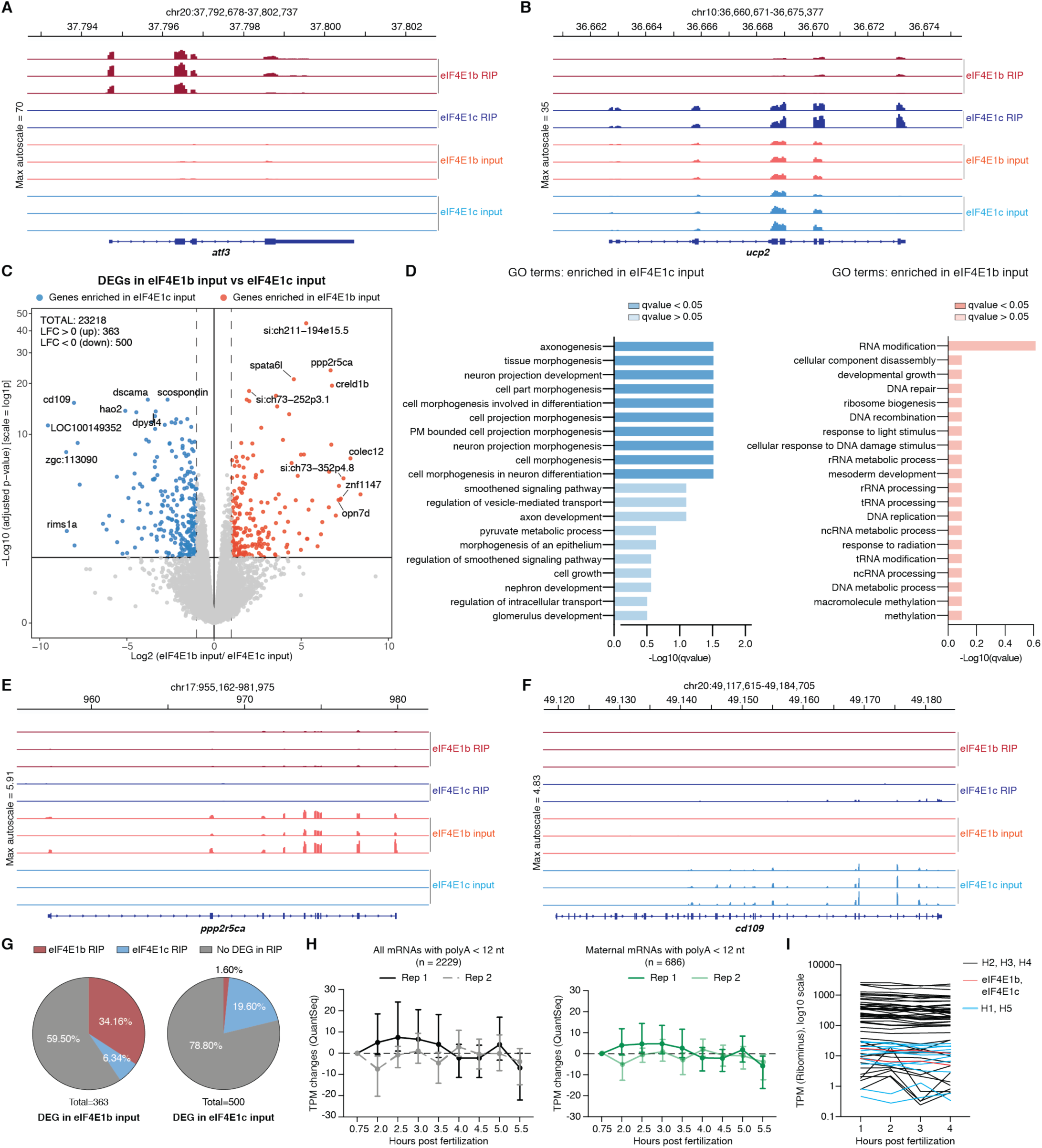
Additional RIP-seq and RNA-seq data analyses. **A-B** Examples of mapped RNA-seq reads of differentially expressed genes (DEGs) in eIF4E1b (*A*) and eIF4E1c (*B*) RNA immunoprecipitations (RIPs). **C** Volcano plot of DEGs in lysates (RIP inputs) from 8-cell embryos expressing 3xflag-sfGFP tagged eIF4E1b or eIF4E1c (p-value < 0.005). **D** Gene ontology (GO) analysis of mRNAs specifically upregulated in embryos overexpressing either eIF4E1c (left) or eIF4E1b (right). **E-F** Mapped RNA-seq reads of example genes upregulated in eIF4E1b (*E*) or eIF4E1c (*F*) inputs. **G** Fraction of transcripts upregulated in embryos overexpressing eIF4E1b (left) or eIF4E1c (right) that are also enriched in eIF4E1b and eIF4E1c RIPs. **H** Changes in mRNA levels (in transcripts per million, TPM) of transcripts with polyA tails containing less than 12 nucleotides during zebrafish embryogenesis according to published RNA-seq data (Bhat *et al*, 2023). All mRNAs are plotted on the left; only mRNAs of maternal origin are plotted on the right. **I** Abundance of histone mRNAs identified in eIF4E1b RIP during the first four hours post fertilization according to published rRNA-depleted RNA-seq data (Cabrera-Quio *et al*, 2021).

## Notes

### Competing Interest Statement

The authors have declared no competing interest.

